# Tissue-resident regulatory T cells exert dualistic anti-tumour and pro-repair function in the exocrine pancreas

**DOI:** 10.1101/2024.02.15.580302

**Authors:** Julie Stockis, Thomas Yip, Shwetha Raghunathan, Celine Garcia, Sheng Lee, Charlotte Simpson, Silvain Pinaud, Martijn J. Schuijs, Joaquín Araos Henríquez, Shaun Png, Gianmarco Raddi, Orian Bricard, Tsz Y. So, Stephanie Mack, Panagiotis Papadopoulos, Ashley Sawle, Duncan I. Jodrell, Andrew N.J. McKenzie, James E.D. Thaventhiran, Menna R. Clatworthy, Giulia Biffi, Kourosh Saeb-Parsy, Adrian Liston, Timotheus Y.F. Halim

## Abstract

Regulatory T cells are fundamentally important for maintaining immune homeostasis, and their potent immune-suppressive roles make them attractive immunotherapeutic targets in cancer. Recent work suggests potential functions of tissue-resident Tregs (trTregs) in tissue-repair and epithelial cell homeostasis. Here, we describe a rare population of trTreg in the exocrine pancreas. We show that these cells share common features of trTregs, including expression of the IL-33 receptor ST2 and production of the epithelial growth factor Amphiregulin, and display an oligoclonal T cell receptor repertoire. Using a mouse model of acute pancreatitis, we show that pancreatic Tregs rapidly expand upon release of IL-33 by fibroblasts. Moreover, depletion of Tregs after initiation of pancreatic injury impairs the regeneration of the exocrine parenchyma. This effect is due, in part, through a direct effect of Tregs on acinar cell proliferation. Finally, we show that transient Treg depletion in established orthotopic pancreatic tumours leads to tumour rejection yet provokes long-lasting damages to surrounding exocrine parenchyma. In all, our results demonstrate the tissue-repair capacity of pancreatic Tregs, and highlight a dualistic role of these cells in the pancreatic tumour ecosystem, with their harmful immune-suppressive function in the tumour coupled to a beneficial tissue-repair function in the surrounding tissue.

## Introduction

Regulatory T cells (Tregs) expressing the transcription factor Foxp3 are indispensable for the prevention of auto-immunity, as exemplified by the uncontrolled immune activation and severe pathologies in Treg-impaired individuals bearing *FOXP3* mutations (Sakaguchi et al., 2020). Importantly, the immunosuppressive capacity of Tregs extends beyond controlling autoimmune diseases, and includes curtailing inflammation after infection, as well as inhibiting CD8^+^ T cell-driven anti-tumour immunity, amongst others. How Tregs suppress immune responses is complex and involves diverse mechanisms, such as the production of the immunosuppressive cytokine transforming growth factor beta (TGF-β) and the inhibition of dendritic cell-mediated T cell activation through the expression of the cell surface molecule cytotoxic T lymphocyte antigen 4 (CTLA-4). Accordingly, Treg-targeted therapies represent an exciting avenue for several diseases, including the immunotherapy of cancer. Notably, antibody-based approaches aiming at depleting Tregs or inhibiting their function, such as anti-CTLA-4 or anti-GARP:latentTGF-β monoclonals, have proven successful in preventing tumour growth in pre-clinical mouse models (de Streel et al., 2020; Simpson et al., 2013). Yet translating these results in the clinic has been challenging, in part due to the occurrence of debilitating immune adverse events.

Emerging studies have revealed the presence of tissue-resident regulatory T cells (trTregs), a distinct Treg subset that populates non-lymphoid tissues (Munoz-Rojas and Mathis, 2021). In addition to immune-suppression, trTregs are critical regulators of organ homeostasis, and/or tissue-repair. For instance, brain trTregs promote neurological recovery following ischemic stroke through the production of the epithelial growth factor amphiregulin (Areg) (Ito et al., 2019), whilst skin trTregs support the hair follicle stem cell niche maintenance through the expression of the cell surface Notch ligand Jagged-1 (Ali et al., 2017). Comparative transcriptomic analysis across a handful of tissues has identified commonalities between trTreg populations over conventional Tregs, such as the expression of *Areg* and *Il1rl1* (encoding a subunit of the IL-33 receptor, ST2), but also suggested potential organ-specific signatures (Munoz-Rojas and Mathis, 2021). These differences likely reveal influence from the tissue niche in instructing the trTreg transcriptome or epigenome. Hence, building a comprehensive descriptive and functional characterisation of trTreg populations in specific organs is of vital importance to understand the consequences of Treg-targeted therapies, as well as to unveil novel tissue-targeted therapies that exploit trTreg function.

In this context, one relatively poorly explored organ is the pancreas, owing to the technical challenges brought by its high content in RNase and other digestive enzymes. Importantly, studies exploring the function of Tregs in the pancreas have mostly focused on their control of anti-islets pathogenic effector T cell responses, and the potential development of Treg cell-based adoptive cell therapy for the treatment of type-1 diabetes (Raffin et al., 2020). In contrast, little is known about the interplay between Tregs and the exocrine parenchyma, which represents the majority of the organ mass. The exocrine pancreas architecture is formed of lobules composed of digestive enzymes-producing acinar epithelial cells structured around a ramified network of ductal cells that are supported by fibroblasts. Interestingly, regeneration of the exocrine pancreas occurs in mice following acute pancreatitis (AP), a common inflammatory injury, which leads to the destruction of the acinar cell compartment (Murtaugh and Keefe, 2015). Clinically, most AP cases will resolve rapidly although rare cases of so-called necrotising pancreatitis can be fatal due to multi-organ failure. Understanding the mechanism underlying pancreatic regeneration has proven difficult. A favoured theory arising from lineage-tracing experiments in mice suggests that regenerated acini originate from spared acinar cells undergoing proliferation (Lodestijn et al., 2021). Whether Tregs contribute to this process has not been investigated.

Here, we sought to identify and comprehensively characterize Tregs of the exocrine pancreas, and assess their role in pancreas regeneration following AP in mice. We found that pancreatic Tregs are uniquely located in the exocrine niche and largely exhibit features of trTregs. Furthermore, we demonstrate that pancreatic Tregs play an indispensable role in tissue repair distinct from their anti-inflammatory role, whether in AP or in pancreatic cancer.

## Results

### trTregs populate the exocrine pancreas

We first questioned whether the pancreas contained trTregs. To do so, we performed flow cytometry on the exocrine pancreas, and assessed the percentage of tissue-resident immune cells by intravenous administration of an anti-CD45 antibody into mice. We observed that the majority of Tregs in the pancreas were unlabelled by the antibody, akin to tissue-resident innate lymphoid cells (ILCs), whilst CD4^+^Foxp3^−^ (Tconv) and CD8^+^ T cells showed increased staining (Figure 1A). Further immunophenotyping analysis by flow cytometry revealed that around half of pancreatic Tregs express the activation marker KLRG1 (Figure 1B), previously shown to mark trTreg population in the skin and visceral adipose tissue (VAT) (Delacher et al., 2017). Little expression of KLRG1 was observed on Tregs isolated from the pancreatic lymph node (pLN), nor on Tconv cells irrespective of location (Figure 1B). Furthermore, KLRG1^+^ pancreatic Tregs expressed high levels of ST2 (Supplementary Figure 1A) and produced higher levels of the epithelial growth factor AREG compared to pLN Tregs (Supplementary Figure 1B), whilst pancreatic KLRG1^−^ Tregs show an intermediate phenotype. Treatment of mice with recombinant IL-33 promoted the expansion of pancreatic Tregs (Figure 1C), as shown previously in other tissues (Kolodin et al., 2015; Schiering et al., 2014; Xu et al., 2018), whether KLRG1^+^ or KLRG1^−^ (Supplementary Figure 1C).

**Figure 1.**
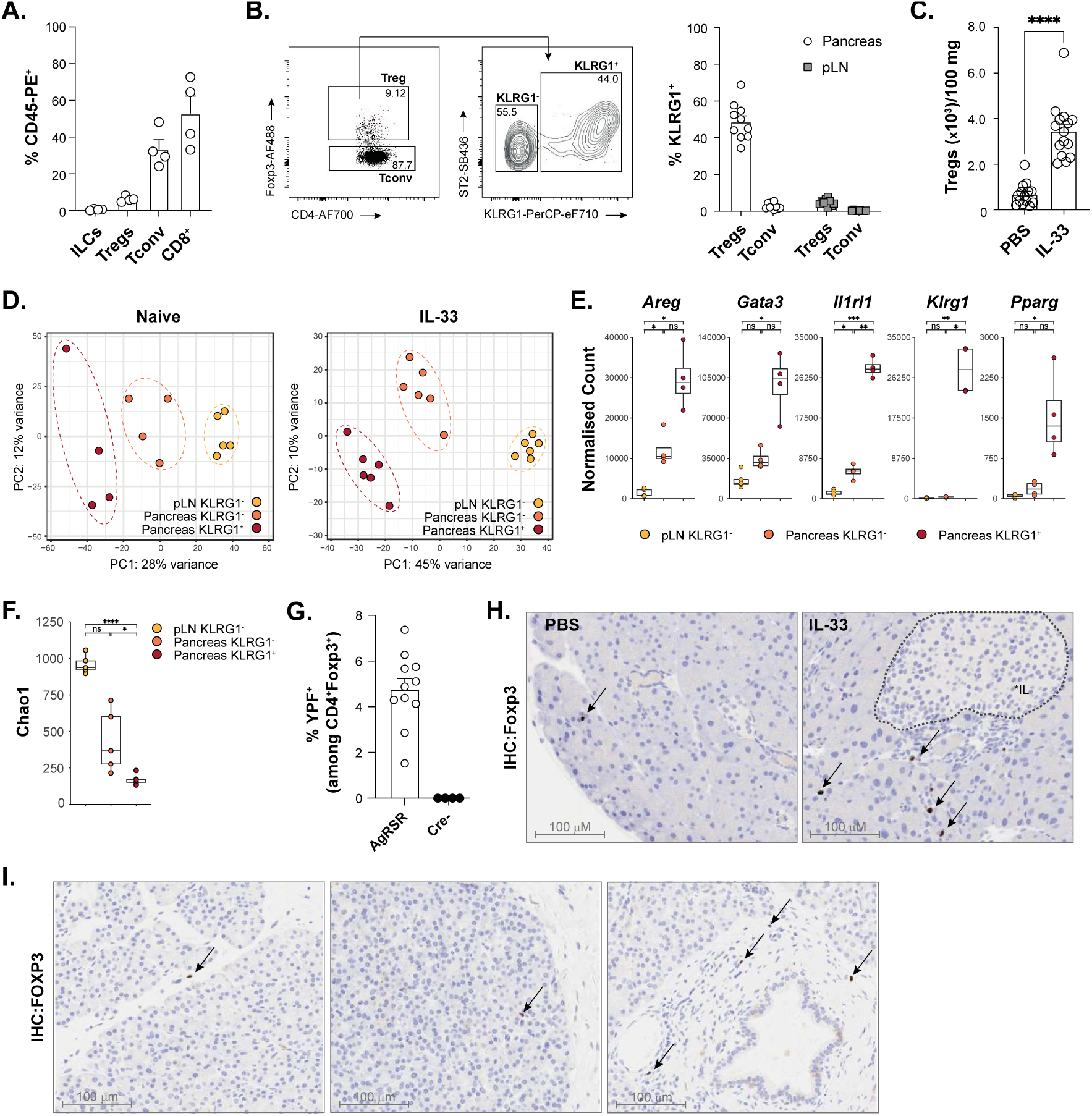
The exocrine pancreas harbours a population of *bona fide* trTregs. Indicated circulating immune cell subsets were quantified in the pancreas of wild-type mice by flow cytometry after i.v. injection of an CD45-PE antibody, followed by sacrifice 5 min later. Results are expressed as %CD45-PE labelled cells among the total indicated cell subset (**A**). Tregs were analysed for KLRG1 expression in the pancreas and pLN of wild-type mice by flow cytometry. Results are shown as a representative dot plot (left) and a bar graph showing the %KLRG1^+^ among Tregs (right) (**B**). Wild-type mice were dosed i.p. with IL-33 on days 0 and 1, and sacrificed on day 5, followed by quantification of total pancreatic Treg numbers by flow cytometry (**C**). Principal component analyses of pancreatic KLRG1^+/−^ and pLN KLRG1^−^ Tregs bulk RNA-seq samples obtained from naïve mice (left, n= 4 to 5 mice) or after IL-33 treatment (right, n= 6 mice, *treated as in C*) (**D**). Expression of the indicated genes in the bulk RNA-seq samples obtained from naïve mice (**E**). TCR-β diversity analysis in the bulk RNA-seq samples obtained from naïve mice (**F**). Mice of the indicated genotype were treated with tamoxifen and sacrificed 7 days later (as indicated in Supplementary Figure 1G), followed by quantification of the number of YFP^+^ cells by flow cytometry (**G**). Wild-type mice were treated as in *C*. Representative images of formalin fixed paraffin embedded (FFPE) sections stained for Foxp3, with black arrows indicating Treg cells (**H**). Representative images of FFPE sections from human healthy pancreas stained for FOXP3, with black arrows indicating Treg cells (**I**). Bar graphs indicate mean (±SEM) and show representative data of 2 independent experiments (n=4 to 5 mice per group, A), or pooled data from 2 to 3 independent experiments (n=4 to 5 mice per group, B, C, G). Histological stainings show representative images of 5 mice per group (H) or 5 human donors (I). * = p ≤ 0.05, ** = p ≤ 0.01, *** = p ≤ 0.001, **** = p ≤ 0.0001, ns = not significant.

To gain further insights into the molecular diversification of pancreatic Treg subsets, we performed bulk RNA sequencing of KLRG1^+^ and KLRG1^−^ Tregs from the pancreata of *Foxp3^YFP^* reporter mice, as well as of pLN KLRG1^−^ Tregs. Given the rarity of Tregs in naïve mice (Figure 1 C), and the high abundance of RNase in the pancreas, we employed a modified version of the SMART-seq2 protocol initially optimised for acinar cell transcriptomics (Wollny et al., 2016). We also separately generated pancreatic Treg transcriptomic data from IL-33-injected animals. Principal component analysis of our two datasets revealed that the transcriptome of pancreatic Tregs differed strongly from that of their pLN counterparts (Figure 1D), with the KLRG1^+^ Tregs segregating furthest from the pLN Tregs, and pancreatic KLRG1^−^Tregs sitting in between. A closer look at canonical trTreg signature genes, including *Areg*, *Gata3*, *Il1rl1*, *Klrg1* and *Pparg*, encoding the master transcription factor peroxisome proliferator-activated receptor (PPAR)-g involved in VAT trTreg function (Cipolletta et al., 2012; Li et al., 2018a), showed a similar pattern as the one observed on the PCA plots, with the highest expression in KLRG1^+^ Tregs compared to pLN, and KLRG1^−^ Tregs showing intermediate levels (Figure 1E). These results are consistent with a stepwise model of trTreg differentiation (Delacher et al., 2020; Li et al., 2018a), but suggest that intermediate populations may also be present within the tissue. Integrated cross-organ transcriptomic analysis using published datasets (Munoz-Rojas and Mathis, 2021) revealed that pancreatic KLRG1^+^ Tregs most closely resemble VAT trTregs, in agreement with their expression of *Pparg*, and differ the most from brain and intestine trTregs (Supplementary Figure 1D).

We further analysed the TCR repertoire of pancreatic Tregs and found that pancreatic Tregs display a much less diverse repertoire than that of pLN Tregs (Figure 1F and Supplementary Figure 1E-F), in agreement with findings in other organs (Munoz-Rojas and Mathis, 2021). To corroborate these findings, we made use of mice bearing a fate-mapper for TCR signalling (*Nr4a1^Kathushka-CreERT2^ Rosa26^LSL-YFP^*, termed AgRSR mice). We found that a single injection of tamoxifen to AgRSR mice resulted in the expression of YFP in 5% of pancreatic Tregs (Figure 1G and Supplementary Figure 1G), consistent with ongoing antigen recognition by trTreg within the pancreatic tissue. Accordingly, expression of the genes encoding members of the Nr4a family were also found to be higher in pancreatic Tregs compared to their pLN counterparts (Supplementary Figure 1H). These results demonstrate a continuous detection of strong TCR signals by pancreatic Tregs that may be driven by the recognition of local presentation of antigens.

Finally, we asked where pancreatic Tregs were located. Immunohistochemistry staining of mouse and human healthy pancreata revealed very few Foxp3-positive cells whilst numbers increased in IL-33 treated mice (Figure 1H and 1I), in agreement with our flow cytometry data (Figure 1C). Interestingly, none of these cells were detected within, or near islets of Langerhans. Instead, we detected Tregs along fibroblast septae, within exocrine acini, and in adventitial cuffs around pancreatic ducts (Figure 1H and 1I). Together, these results demonstrate that the pancreas harbours a population of bona-fide trTregs in the exocrine parenchyma.

### Treg expand during pancreatic inflammation

The unexpected location of Tregs in the exocrine, but not endocrine, pancreas begged the question of whether crosstalk between Tregs and the exocrine niche exists. Injection of the cholecystokinin orthologue caerulein induces AP, and provides a well-established model for the study of tissue repair after acute injury to the exocrine parenchyma. We treated mice with 6 hourly injections of caerulein to trigger AP, and observed that within 24 hours, pancreatic acini showed regions of necrosis, oedema with enlarged septae, and infiltration of immune cells (Figure 2A). By day 3 many acinar cells displayed a ductal morphology via a protective process known as acinar-to-ductal metaplasia, while by day 7 the parenchyma had returned to a normal architecture (Figure 2A). We queried whether Tregs expand during this process, and found that Tregs increased in number at day 3 following pancreatic injury (Figure 2B). Interestingly, we observed a rapid induction of pancreatic Treg cycling at day 1 post-injury (Figure 2C), before we could detect increases in their absolute numbers (Figure 2B); this suggests a contribution from local expansion of trTregs to the increased numbers seen on day 3. Furthermore, we found that Treg proliferation was reduced in mice deficient in IL-33 (Figure 2C). This was not the case when preventing antigen recognition using an anti-MHC-II blocking antibody (Supplementary Figure 2A and 2B). Thus, these results suggest that pancreatic trTreg sense tissue damage via the release of IL-33, which contributes to the accumulation of Tregs seen in AP in a TCR-independent manner.

**Figure 2.**
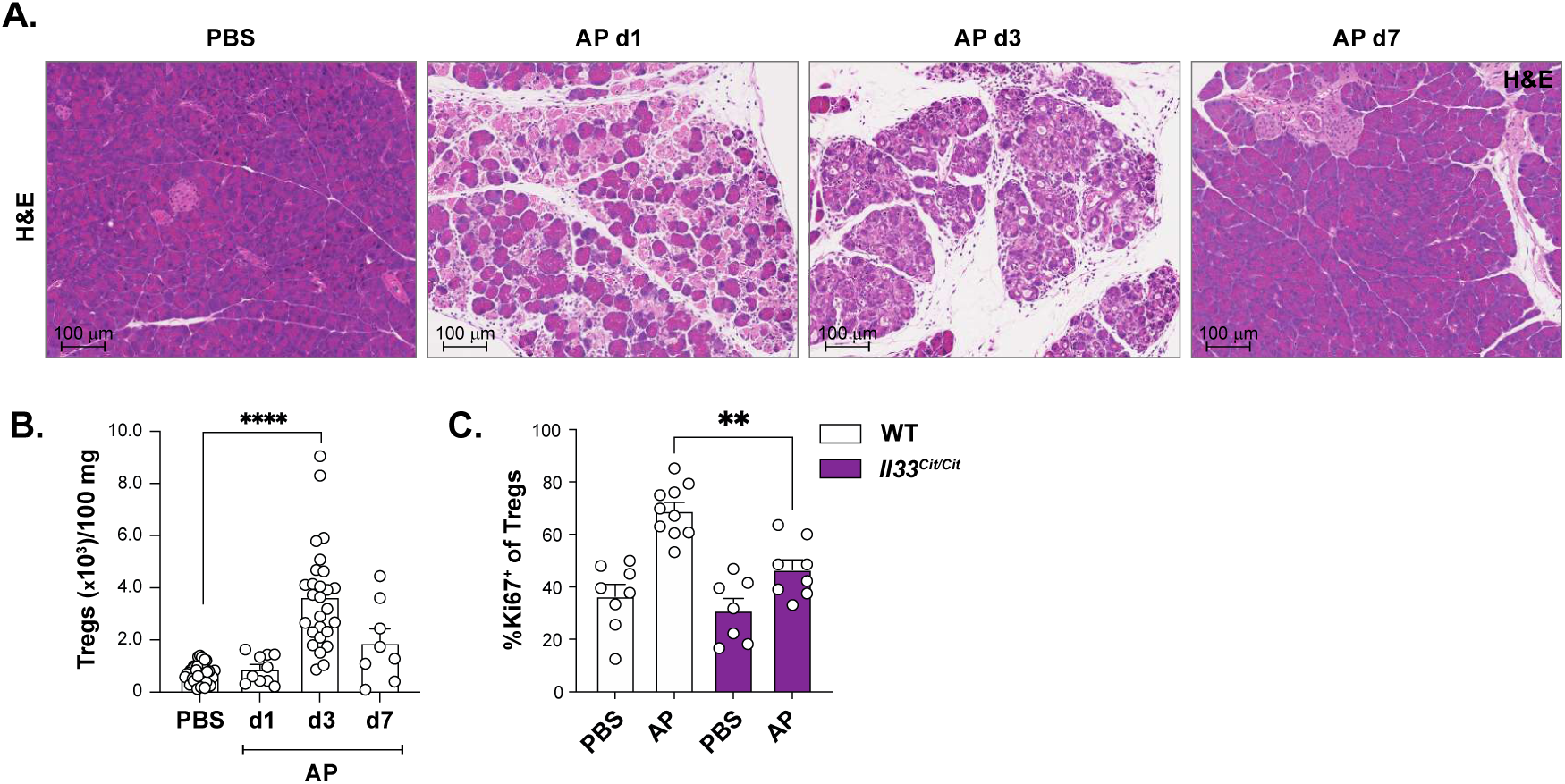
Treg expansion during pancreatic inflammation is IL-33-dependent. Mouse pancreas histology at different timepoints after induction of acute pancreatitis (AP). Representative haematoxylin and eosin (H&E) stained FFPE sections representing steady-state (PBS), acinar cell necrosis and acute injury (day 1), acinar-to-ductal metaplasia (day 3), and resolution of tissue architecture (day 7) (**A**). Wild-type mice were sacrificed at different timepoints after induction of AP, followed by quantification of total pancreatic Treg numbers by flow cytometry (**B**). Mice of the indicated genotype were sacrificed one day after induction of AP, followed by quantification of cycling Tregs by flow cytometry (**C**). Bar graphs indicate mean (±SEM) and show pooled data from 2 (n=3 to 5 mice per group, C) to 11 independent experiments (n=3 to 5 mice per group, B). * = p ≤ 0.05, ** = p ≤ 0.01, *** = p ≤ 0.001, **** = p ≤ 0.0001, ns = not significant.

### Fibroblasts, but not acinar cells, are the source of IL-33 in the pancreas

It was previously reported that acinar cells are the source of IL-33 during pancreatitis (Kempuraj et al., 2013; Watanabe et al., 2016), although recent studies have implicated fibroblasts as an important source of IL-33 responsible for the maintenance and expansion of trTregs and type-2 ILCs (ILC2) (Mahlakoiv et al., 2019; Spallanzani et al., 2019). We used flow cytometry to identify major non-immune populations in the pancreas (Supplementary Figure 3A, left panel). We subsequently examined *Il33* gene expression in sorted acinar cells, ductal cells and fibroblasts from naïve mice and found that only fibroblasts expressed *Il33* (Figure 3A). Furthermore, 2-photon imaging of *ex vivo* pancreata from *Il33^Cit/+^* reporter mice showed co-localization of citrine fluorescence with collagen fibres (Supplementary Figure 3B). Finally, antibody-mediated staining of IL-33-producing cells in pancreas sections from naïve mice confirmed their mesenchymal origin, whilst no acinar cells were found to express IL-33 (Figure 3B).

**Figure 3.**
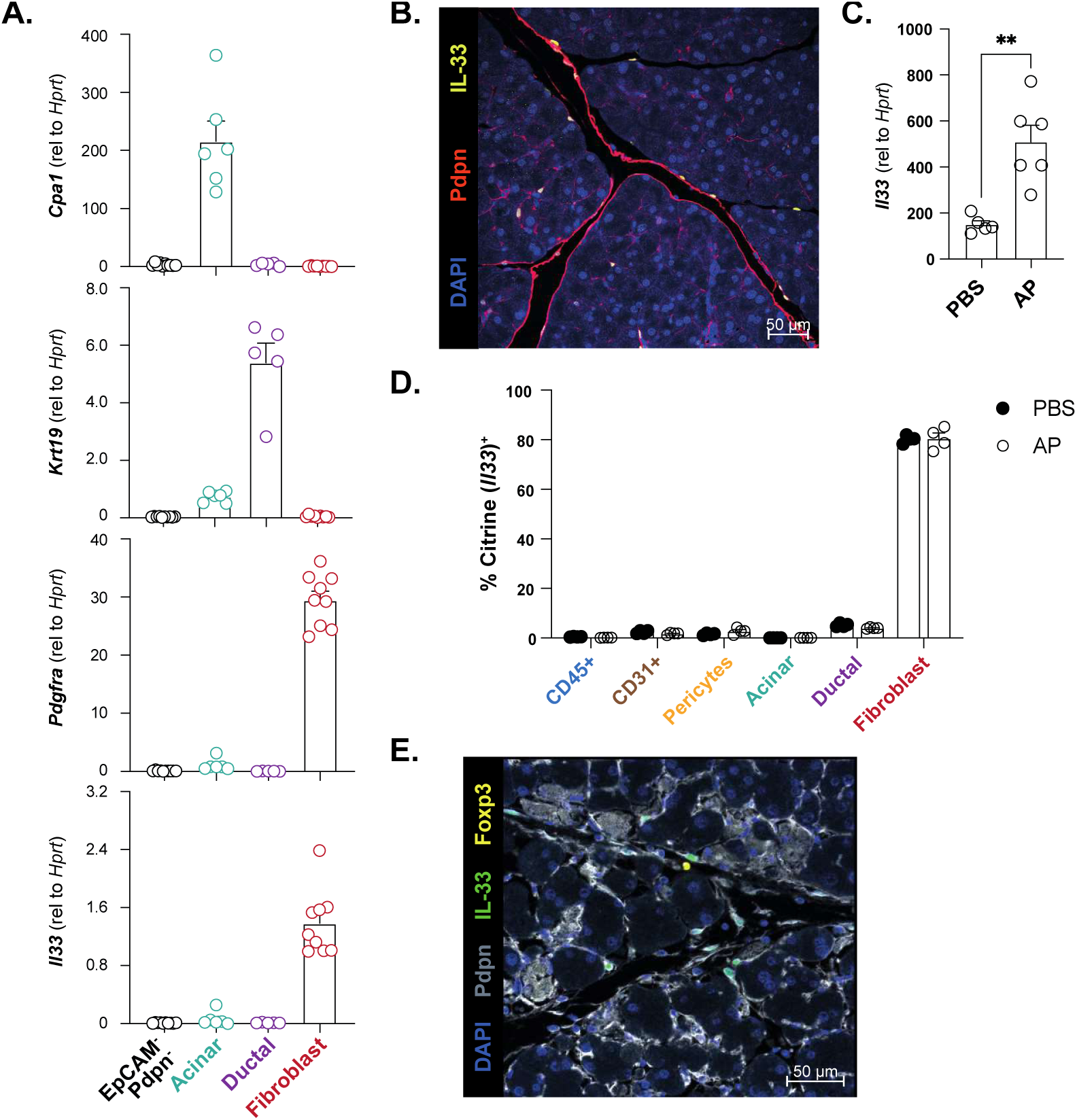
Fibroblasts, but not acinar cells, are the source of IL-33 in the pancreas. Pancreatic non-immune cells from wild-type mice were sorted by FACS, followed by RNA extraction and RT-qPCR for the indicated genes. Acinar cells were gated as CD45^−^ EpCAM^+^CD133^lo^SSC^hi^, ductal cells as CD45^−^EpCAM^+^CD133^+^SSC^lo^, fibroblasts as CD45^−^ EpCAM^−^Pdpn^+^. Remaining non-immune cells were sorted as CD45^−^EpCAM^−^Pdpn^−^ and contain mostly endothelial cells (**A**). FFPE pancreata were sectioned and analysed using immunofluorescence (IF) imaging. A representative region is shown and indicates co-localisation of Podoplanin (Pdpn) and IL-33 staining (**B**). Wild-type mice were sacrificed 6 hours after induction of AP. Total pancreata were harvested for RNA extraction, followed by RT-qPCR for *Il33* (**C**). *Il33^Cit/+^* or control mice were sacrificed 6 hours after induction of AP, and analysed by flow cytometry to determine the phenotype of IL-33^Cit+^ cells. Gating strategy is depicted in Supplementary Figure 3A (**D**). Multiplex IF imaging of FFPE sections from wild-type mice 24 hours after induction of AP. A representative region is shown and indicates proximity between IL-33 and Foxp3 staining (**E**). Bar graphs indicate mean (±SEM) and show representative data of 1 (n=3 to 4 mice per group, C) or 2 independent experiments (n=4 mice per group, D), or pooled data from 2 independent experiments (n=5 mice per group, A). (m)IF stainings show representative images of 2 to 3 mice analysed (B, E). * = p ≤ 0.05, ** = p ≤ 0.01, *** = p ≤ 0.001, **** = p ≤ 0.0001, ns = not significant.

Interestingly, we observed that *Il33* gene expression was already increased in total pancreata 6 hours after injury (Figure 3C), supporting our data showing IL-33-dependent Treg cycling early after damage (Figure 2C). Using *Il33^Cit/+^* reporter mice, we further showed that the only cells expressing *Il33* during AP were fibroblasts (Figure 3D and Supplementary Figure 3A, middle and right panel), similar to our observations in naïve animals. Finally, we observed that pancreatic Tregs are localised in the vicinity of IL-33-producing fibroblasts at 24h post-injury (Figure 3E). Thus, our findings demonstrate that IL-33 is produced by fibroblasts, but not acinar cells, in agreement with a recent report describing IL-33^+^ fibroblasts in chronic pancreatitis (Yang et al., 2022), providing for release of IL-33 during AP.

### Tregs play a non-redundant role in pancreas regeneration

We asked whether Tregs played a functional role at different stages during pancreatic injury. We first assessed whether Tregs were important for the initial damage phase of AP. To this end, we depleted Tregs before the start of injury using *Foxp3^DTR^* mice in which expression of the diphtheria toxin receptor (DTR) is driven from the endogenous *Foxp3* promoter specifically in Tregs (Figure 4A). Treatment of mice with diphtheria toxin (DT) promoted efficient Treg depletion (Supplementary Figure 4A) yet did not result in any measurable difference in the severity of AP at 24h compared to control mice, as assessed by serum amylase levels and CD45^+^ immune cell infiltration (Figure 4B). To assess whether Treg played a role in pancreas regeneration during AP, we instead depleted Tregs after the initiation of injury (Figure 4C). Examination of the pancreas architecture at day 7 following injury revealed a striking destruction of the exocrine parenchyma in Treg-depleted mice compared to control mice (either wild-type mice treated with DT, or *Foxp3^DTR^*mice littermate controls) along with immune infiltration and metaplastic cells characterised by the expression of the ductal marker cytokeratin 19 (CK19) (Figure 4D). Flow cytometry analyses confirmed the depletion of Tregs along with a 10-fold increase in total immune cell numbers (Supplementary Figure 4B). Altogether, these results demonstrate that Treg depletion does not impede tissue damage onset but impairs regeneration following injury.

**Figure 4.**
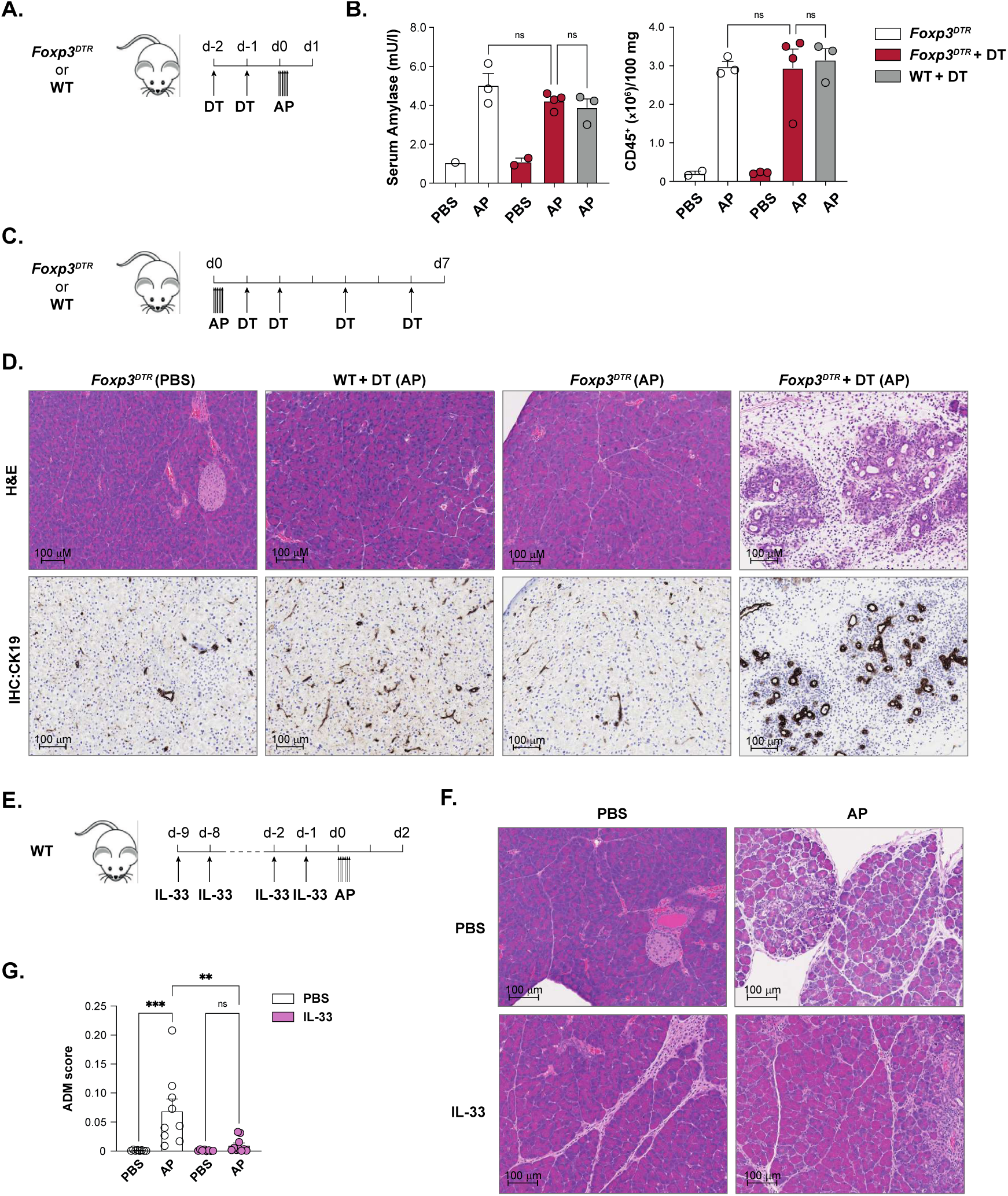
Tregs play a non-redundant role in pancreas regeneration. A-B. Mice of the indicated genotype were treated with diphteria toxin (DT) before induction of AP and sacrificed one day later (**A**), followed by quantification of serum amylase levels (**B**, left panel) and total CD45^+^ numbers by flow cytometry (**B**, right panel). **C-D.** Mice of the indicated genotype were treated with DT after the induction of AP and sacrificed 7 days later (**C**). Representative H&E (top) and CK19 (bottom) stained FFPE sections are shown (**D**). **E-G.** Wild-type mice were dosed with IL-33 before induction of AP and sacrificed 2 days later (**E**). Representative H&E stained FFPE sections are shown (**F**). ADM scoring on H&E stained FFPE sections was performed with the HALO image analysis software (**G**). Bar graphs indicate mean (±SEM) and show representative data of 2 independent experiments (n=2 to 4 mice per group, B), or pooled data from 2 independent experiments (n=3 to 5 mice per group, G). * = p ≤ 0.05, ** = p ≤ 0.01, *** = p ≤ 0.001, **** = p ≤ 0.0001, ns = not significant.

We next asked whether we could increase Treg numbers and hence accelerate the regeneration process. As we showed that IL-33 efficiently expanded pancreatic Tregs (Figure 1C), we repeatedly dosed wild-type mice with IL-33 before the initiation of injury (Figure 4E) and sacrificed the mice two days later. Flow cytometry analysis confirmed that dosing of mice with IL-33 expanded pancreatic Tregs, but did not influence the total number of CD45^+^ cells after AP (Supplementary Figure 4C). Moreover, whilst control mice showed typical features of ADM two days after injury, mice pre-dosed with IL-33 showed an almost intact parenchyma, albeit with signs of immune infiltration (Figure 4F). Quantification of ADM lesions using a machine learning algorithm confirmed these observations (Figure 4G and Supplementary Figure 4D). Thus, these results suggest that increased Treg numbers are associated with improved outcome during AP. A caveat to this experiment lies in the IL-33-driven expansion of immune cells other than Tregs, most notably ILC2s. Yet, we and others have shown that ILC2s play a detrimental role in pancreatitis (Yip *et al*., submitted)(Yang et al., 2022). Thus, the protective effect observed in AP upon IL-33 pre-treatment is unlikely to be mediated by ILC2s. Furthermore, repeating this experiment in Treg-deficient animals is however not possible given the auto-immunity that would arise from a 2-week long depletion regimen.

In all, our data support a non-redundant role for Treg cells in pancreas regeneration following acute injury.

### Tregs promote acinar cell proliferation directly

The mechanisms underlying exocrine pancreas regeneration are not fully understood; one hypothesis suggests that mature acinar cells can act as facultative stem cells after injury, and that their proliferation is required for subsequent regeneration (Desai et al., 2007; Wollny et al., 2016). Using our non-immune flow cytometry panel and Ki67 staining, we indeed found acinar cells cycling at day 7 post-injury, whilst these were almost absent in naïve PBS-treated animals (Figure 5A and 5B). Strikingly, acinar cell proliferation was substantially reduced in Treg-depleted mice (Figure 5A and 5B), correlating with an impaired regeneration (Figure 4D). This could not be attributed to a toxic effect of DT, as wild-type mice subjected to pancreatitis followed by DT treatment still show significantly more acinar cell cycling than Treg-depleted animals (Figure 5B). Importantly as well, this effect was specific of acinar cells, as Treg depletion had no significant effect on ductal cell cycling (Figure 5C).

**Figure 5.**
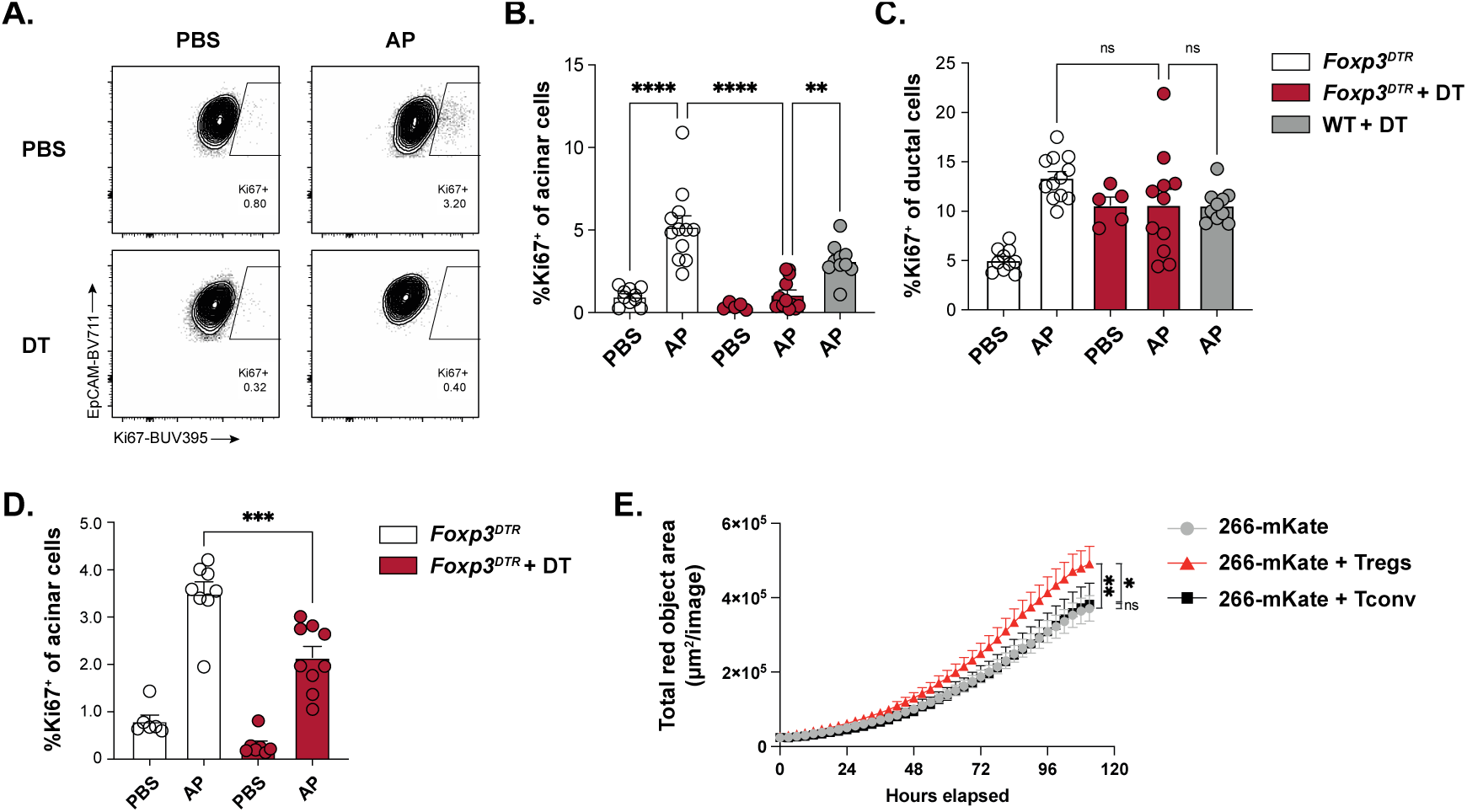
Treg promote pancreas regeneration through a direct effect on acinar cell proliferation. A-C. Mice of the indicated genotype were treated with DT after the induction of AP and sacrificed 3 (**D**) or 7 days (**A-C**) later. Representative dot plot (**A**) and quantification of cycling acinar (**B**, **D**) and ductal cells (**C**) by flow cytometry. Sorted pancreatic Tregs or Tconv cells were co-cultured with 266-6 acinar cells expressing the red fluorescent protein m-Kate in the presence of anti-CD3/anti-CD28 coated beads, IL-2 and IL-33. Acinar cell proliferation was measured in the Incucyte live imaging system (**D**). Bar graphs indicate mean (±SEM) and show pooled data from 2 to 3 independent experiments (n=3 to 5 mice per group, B-D). Growth curve shows data representative of 2 independent experiments. * = p ≤ 0.05, ** = p ≤ 0.01, *** = p ≤ 0.001, **** = p ≤ 0.0001, ns = not significant.

Given that trTregs can promote tissue repair via direct effects on epithelial cells, we asked whether the decrease in acinar cell proliferation and associated lack of regeneration following Treg depletion could be the result – at least in part – of a direct effect of Tregs on acinar cells, or alternatively, whether it would merely be the consequence of the heightened inflammation triggered by Treg loss. In support of the former hypothesis, we first found that Treg depletion already impairs acinar cell proliferation at day 3 post-injury (Figure 5D), a time point where Treg depletion has not yet impacted immune cell numbers in the pancreas (Supplementary Figure 5A-B) thereby excluding the recruitment of inflammatory cells as a potential cause for reduced acinar cell cycling. Secondly, we made use of *Foxp3^DTR/x^* heterozygous females, wherein half of the Treg express DTR owing to the random inactivation of the X chromosome, which harbours the *Foxp3* locus. Thus, in these mice, treatment with DT only allows for 50% Treg depletion, leaving the remaining 50% Tregs capable of preventing overt inflammation and auto-immune manifestations. Interestingly, examination of *Foxp3^DTR/x^* pancreata at day 7 post-injury revealed an increased trend, although not significant, in parenchymal area showing remaining ADM lesions (Supplementary Figure 5C-E). Finally, we tested *in vitro* whether Treg could directly promote acinar cell proliferation. As primary acinar cell cultures are challenging and proved difficult to robustly control, we turned to the 266-6 acinar cell line, which we transduced with a lentivirus encoding a red reporter (266-6-mKate) to allow for live cell imaging. We further sorted pancreatic Tregs from *Foxp3^YFP^* reporter mice and co-cultured them with the 266-6-mKate cells (Supplementary Figure 5F). Here, we showed that pancreatic Tregs, but not CD4^+^Foxp3^−^ T cells (Tconv), were able to directly enhance the growth of 266-6 cells (Figure 5E). Altogether, these results support the notion that Tregs can directly promote acinar cell proliferation after acute damage to the exocrine pancreas.

### A Treg-acinar cell interaction network

To uncover potential mechanisms whereby pancreatic Tregs support the proliferation of acinar cells upon tissue injury, we examined our bulk RNA-seq dataset generated from IL-33 treated animals. Given that KLRG1^+^ pancreatic Tregs displayed the most mature, effector phenotype (Figure 1D-F), we selected genes that were significantly upregulated in this population compared to pancreatic KLRG1^−^ and pLN Tregs, yielding 1186 genes (FDR<0.01) (Figure 6A, yellow line). We further refined this list by selecting genes that were also upregulated in pancreatic KLRG1^−^ Tregs compared to pLN, leaving us with 601 genes (FDR<0.01) (Figure 6A, red line). Among others, the resulting gene list contained canonical pan-trTreg genes such as *Areg*, *Gata3* and *Il1rl1* (Supplementary Figure 6A and Supplemental Table 1), showing a similar expression pattern as the one observed in naïve mice (Figure 1E).

**Figure 6.**
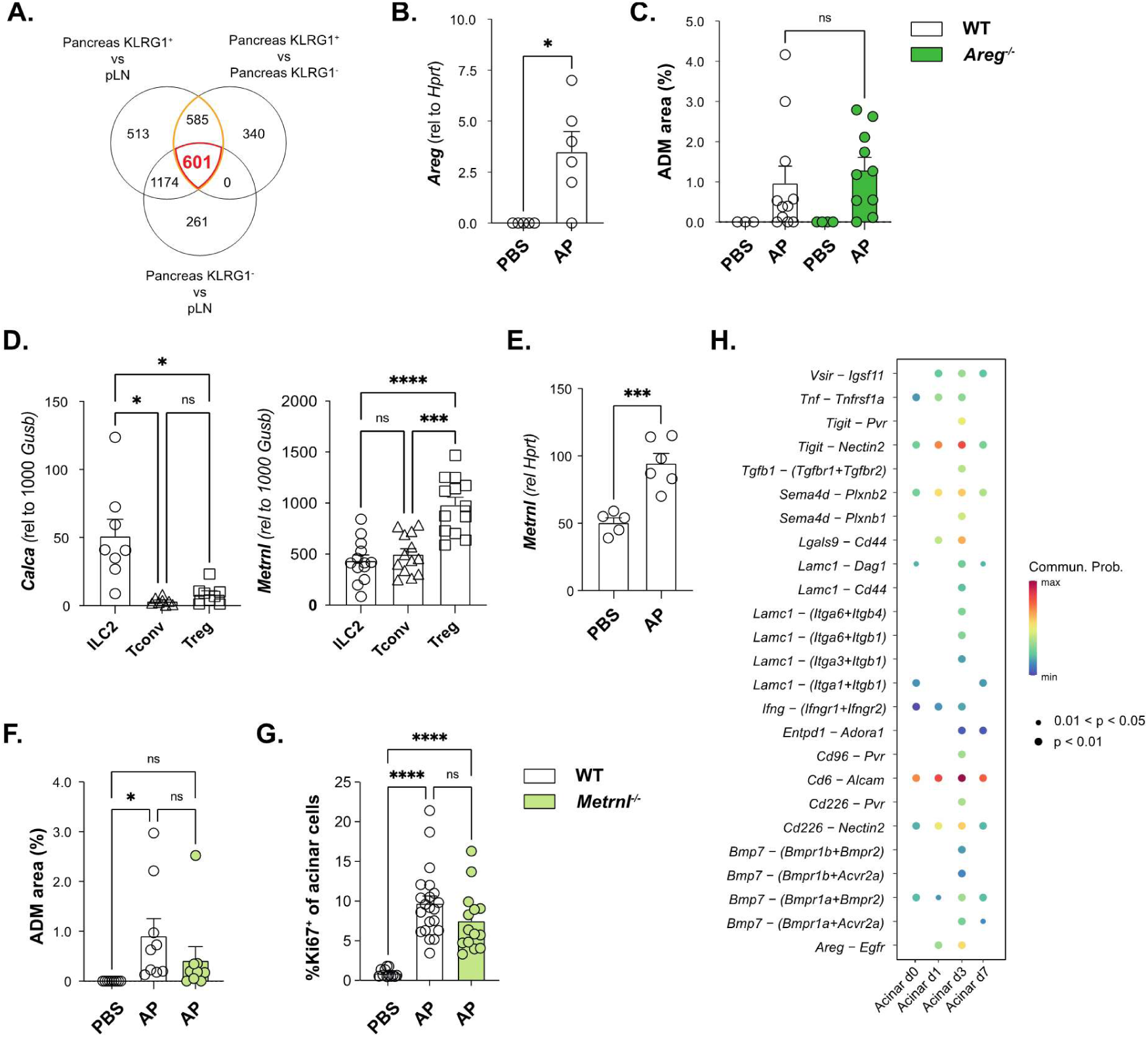
Crosstalk between pancreatic Tregs and acinar cells. Venn diagrams showing the number of genes significantly upregulated in the indicated comparisons obtained from the differential gene expression analysis of the bulk RNA-seq dataset from IL-33 treated mice (as described in Figure 1D, right) (**A**). Wild-type mice were sacrificed 6 hours after induction of AP. Total pancreata were harvested for RNA extraction, followed by RT-qPCR for *Areg* (**B**) and *Metrnl* (**E**). Mice of the indicated genotype were sacrificed 7 days after induction of AP, followed by quantification of ADM area on H&E sections using the HALO software (**C**). ILC2s, Tconv and Tregs were FACS-sorted from wild-type mice treated with IL-33, followed by RNA extraction and RT-qPCR for *Calca* (left) and *Metrnl* (right) (**D**). Mice of the indicated genotype were sacrificed 7 days after induction of AP, followed by quantification of ADM area on H&E sections using the HALO software (**F**) and of cycling acinar cells by flow cytometry (**G**). Treg to acinar cells ligand-receptor pairs were inferred with the Cell-Chat tool, using single-cell transcriptomic datasets from pancreatic trTregs and a time course of acinar cells after AP (**H**). Bar graphs indicate mean (±SEM) and show representative data of 1 independent experiments (n=3 to 4 mice per group, B, E), or pooled data from 2 (n=3 to 6 mice per group, C, F), 3 (n=3 to 5 mice per group, D) or 4 independent experiments (n=2 to 8 mice per group, G). * = p ≤ 0.05, ** = p ≤ 0.01, *** = p ≤ 0.001, **** = p ≤ 0.0001, ns = not significant.

Treg production of Amphiregulin was previously shown to contribute to tissue repair in the lung, brain and muscle (Arpaia et al., 2015; Burzyn et al., 2013; Ito et al., 2019). Interestingly, *Areg* expression was increased early following pancreatic injury (Figure 6B), and recombinant Areg was able to induce the proliferation of 266-6-mKate acinar cells *in vitro* (Supplementary Figure 6B). However, histological analysis of pancreata of wild-type and *Areg^−/−^* mice were undistinguishable at day-7 post-injury (Figure 6C and Supplementary Figure 6C), arguing against a substantial role for Treg-produced Areg in promoting pancreas regeneration.

To find additional candidates, we restricted the 601-gene list to membrane-bound or secreted candidates, yielding 90 genes (Supplementary Table 2). We then selected candidates linked with regeneration processes in the literature. We found *Bmp7* and *Calca,* encoding ligands with potential roles in liver regeneration (Laschinger et al., 2020; Sugimoto et al., 2007), and *Metrnl*, encoding the cytokine meteorin-like recently implicated in muscle regeneration (Baht et al., 2020). We further assessed the expression pattern of these genes in our RNA-seq dataset from naïve mice, and found that *Bmp7* was not significantly differentially expressed between pancreatic Tregs and their LN counterparts, whilst *Calca* and *Metrnl* were expressed at higher levels in KLRG1^+^ pancreatic Tregs compared to pLN KLRG1^−^ Tregs (Supplementary Figure 6D). However, qPCR analysis on sorted ILC2, Tconv and Treg cells from IL-33-treated animals showed that *Metrnl* expression, but not *Calca*, was enriched in Tregs (Figure 6D). Moreover, expression of *Metrnl* was rapidly increased upon pancreatic injury in total tissue extracts (Figure 6E). In order to determine whether meteorin-like was required for pancreatic regeneration following AP, we generated *Metrnl* whole-body knock-out mice through CRISPR-Cas9 technology. Metrnl-deficient animals were viable and born at mendelian ratio, without any gross functional abnormalities, as described previously (Baht et al., 2020). However, histological examination did not reveal any significant defect in regeneration 7 days post-injury in *Metrnl^−/−^* animals compared to control animals (Figure 6F). Accordingly, we could not detect any significant difference in acinar cell cycling by flow cytometry, although there was a trend towards a decreased acinar cell proliferation in *Metrnl^−/−^* animals compared to control mice (Figure 6G).

We separately performed scRNA-seq to generate a dynamic atlas of non-immune cell types over the course of AP (Yip *et al*., submitted). We used this dataset along with a pre-published scRNA-seq dataset of pancreatic Tregs (Burton et al., 2023) to interrogate unbiasedly ligand-receptor interactions between acinar cells and pancreatic Tregs at different timepoints post injury. Besides *Bmp7* and *Areg*, CellChat analysis revealed other signalling pathways involved in epithelial repair, namely the adenosine pathway (*Entpd1*), the semaphorin-plexin pathway (*Sema4d*), and the production of Galectin-9 (*Lgals9*) (Figure 6H). Moreover, this analysis also predicted that Tregs could engage with acinar cells via laminin-integrin interactions (*Lamc1*), as well as via immune checkpoint molecules (*Vsir*, *Tigit*, *Cd96*, *Cd206*), which may contribute to the retention of Tregs within the exocrine parenchyma.

Collectively, our data suggest that pancreatic Tregs can engage with acinar cells via multiple signalling networks, some of which may be relevant for the pro-proliferative effect of Tregs on acinar cells and warrant further study.

### Treg depletion promote pancreatic tumour rejection yet leads to pancreatitis

Finally, we asked whether the crucial role played by Tregs in pancreas regeneration was affected by Treg targeting in pancreatic cancer. As shown by others (Clark et al., 2007), we found that Tregs are elevated in pancreatic neoplastic lesions of *Ptf1a^Cre/+^Kras^LSL-G12D/+^* (KC) mice (Figure 7A). Furthermore, Treg depletion after orthotopic implantation of a *Pdx1^Cre/+^Kras^LSL-G12D/+^Trp53 ^LSL-R172H/+^Rosa26 ^YFP/YFP^* (KPCY)-derived pancreatic tumour cell line dramatically improved survival (Figure 7B and Supplementary Figure 7A). Surprisingly, histological analysis of pancreata at study endpoint revealed some pancreatitis-like area in the Treg-depleted group, suggesting unresolved ADM (Supplementary Figure 7B). To build on this observation, we further analysed pancreata of Treg-depleted and control animals at 21 days post-tumour implantation. In control animals, tumour lesions were already visible and well constrained in a stromal capsule, whilst the exocrine parenchyma outside of the tumour margin was mostly unremarkable (Figure 7C and Supplementary Figure 7C). In contrast, Treg-depleted animals were already devoid of histologically identifiable tumour, yet most of the exocrine parenchyma showed ADM-like structures (Figure 7C and Supplementary Figure 7C). Notably, Treg numbers had already recovered to normal in the pancreas of Treg-depleted animals at day 21 (Figure 7D). Thus, our results demonstrate that the non-redundant function of pancreatic Tregs in tissue repair is coupled to their immune-suppressive properties in the tumour ecosystem. As a result, punctual Treg ablation to treat pancreatic cancer results in a pancreatitis-like disease, which can potentially be long-lasting.

**Figure 7.**
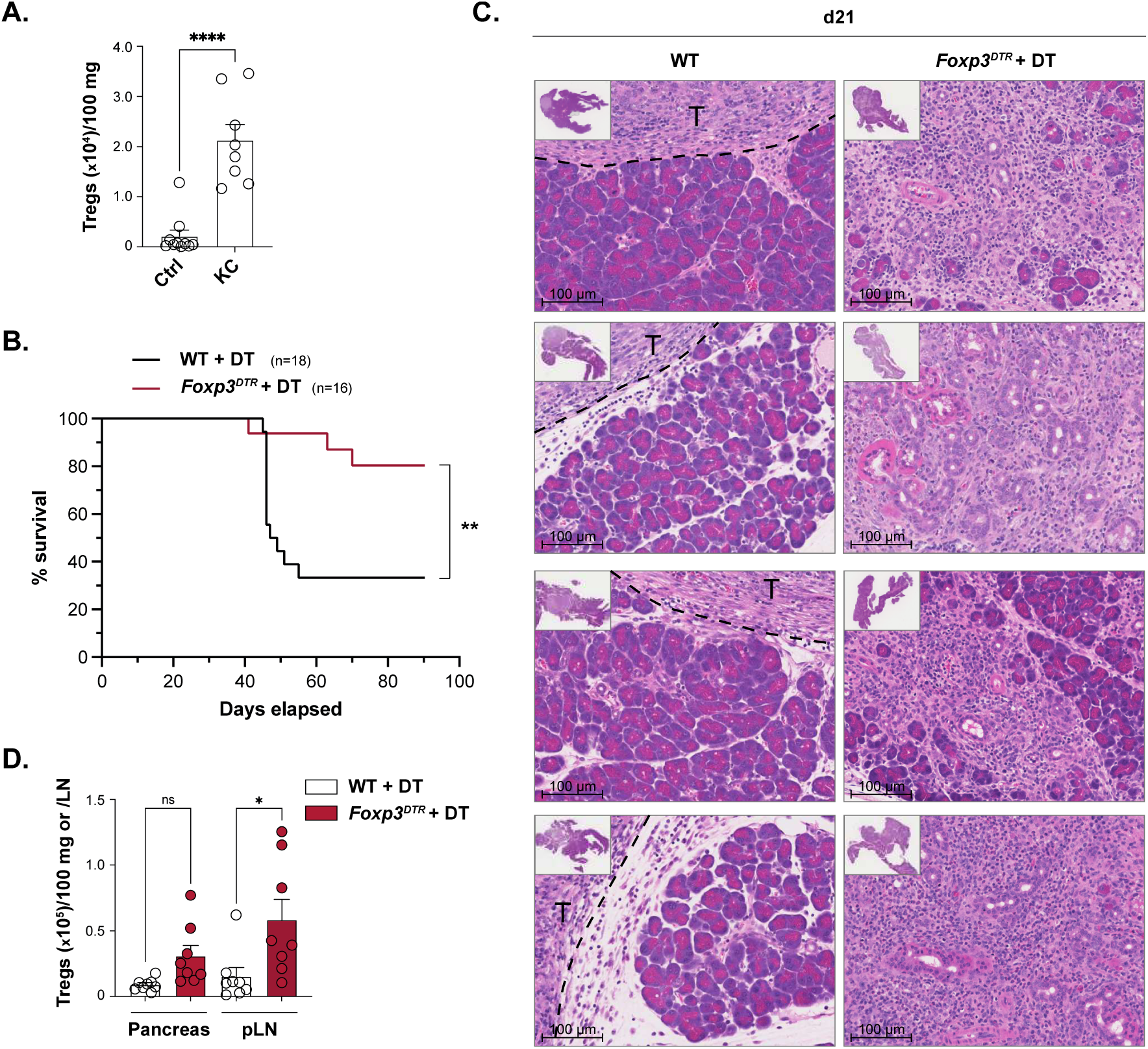
Anti-tumour efficacy of Treg depletion in an orthotopic PDAC model is associated with alterations to the exocrine parenchyma. Pancreatic Tregs from 10-week- to 11-month-old KC mice and littermate controls were quantified by flow cytometry (**A**). 2838c3 cells were implanted orthotopically in the pancreas of mice of the indicated genotype, followed by treatment with DT (as indicated in Supplementary Figure 7A), followed by survival analysis (**B**). Representative H&E sections of mice treated as in *(B)* and sacrificed at 21 days post-implantation **(C)**. Treg numbers were quantified by flow cytometry in the pancreas and pLN of mice treated as in *(B)* and sacrificed at 21 days post-implantation **(D).** Bar graph indicate mean (±SEM) and show representative data of 1 independent experiment (n=8 mice per group, D), or pooled data from 3 independent experiments (n=2 to 4 mice per group, A). * = p ≤ 0.05, ** = p ≤ 0.01, *** = p ≤ 0.001, **** = p ≤ 0.0001, ns = not significant.

## Discussion

It is increasingly clear that Tregs maintain organ homeostasis not only by controlling the function of other immune cells but also by actively engaging with non-immune cells in their surrounding tissue niche. In the pancreas, Tregs locally enforce control over NK cell-derived IFNγ to prevent the destruction of endocrine beta cells by pathogenic T cells (Feuerer et al., 2009), and over pro-fibrotic type-2 innate and adaptive immune cells in the context of chronic pancreatitis (Glaubitz et al., 2022). Whether Tregs directly contribute to tissue repair in this organ had not been investigated. Here we show that Tregs play an essential role in mediating pancreas repair following acute injury. We found that release of IL-33 from fibroblasts during AP promotes Treg expansion; moreover, trTregs infiltrate the exocrine parenchyma and assist acinar cell proliferation to ensure tissue regeneration.

Mechanistically, our data support a direct effect of trTreg-derived signals in instructing acinar cell proliferation, although we could not pinpoint the precise identity of such signals. Interestingly, our observation is consistent with the reported pro-proliferative effect of Tregs on epithelial stem cells from the hair follicle upon hair regeneration (Ali et al., 2017). In that case, comparison of skin Tregs and their lymphoid counterparts pointed towards the Notch ligand Jagged-1 as underpinning this effect. Accordingly, we found that pancreatic Tregs express *Jag1* (Supplemental Table 1) although we could not confirm its surface expression (data not shown). Given the impaired pancreas regeneration observed in Notch1-deficient mice upon injury (Siveke et al., 2008), it would be interesting to determine whether Treg-specific Jag-1-deficient mice display a similar phenotype. Other pancreatic Treg-derived signals uncovered by our transcriptomic analysis included amphiregulin and the novel cytokine meteorin-like; strikingly, both *Areg* and *Metrnl* expression were increased quickly after pancreatic injury. However, we could not find compelling *in vivo* evidence for a role of these secreted factors in pancreas regeneration. Lastly, our unbiased computational network analysis revealed a putative ligand-receptor interactome between Treg and acinar cells over the course of acute injury, opening new avenues for the exploration of these pairs in Treg-mediated pancreas regeneration.

An open question arising from our study concerns the nature of the antigen recognised by pancreatic trTregs. Here we found that pancreatic Tregs display an oligoclonal TCR repertoire, in line with other studies on trTregs (Munoz-Rojas and Mathis, 2021). Using AgRSR mice, we further provide evidence for the recognition of local antigens by pancreatic Tregs. In a mouse model of autoimmune diabetes, pancreatic Treg TCRs were found to react against islet-derived antigens, amongst which insulin and proinsulin (Spence et al., 2018). In this context, however, Tregs were isolated from the inflamed islets of the endocrine pancreas. Herein we show that in naïve mice as well as in humans, pancreatic Tregs populate the exocrine parenchyma or the fibroblast-rich interstitial septae and were not inhabiting the endocrine islets. Thus, we speculate that a large proportion of pancreatic Tregs recognise yet unidentified exocrine parenchymal cell-derived antigens. Availability of paired αβ TCR sequences from scRNA-seq data is now enabling TCR reconstruction and screening for their cognate antigen in reporter assays; such approaches will undoubtedly reveal new determinants of trTreg function and ontogeny.

A detailed picture of the non-immune functions exerted by trTreg populations across distinct tissues in homeostasis and upon injury is now emerging. This compendium bears particular relevance for the development and use of Treg-targeted therapies in various disease contexts, including the immunotherapy of cancer. In the pancreas, genetic or antibody-mediated Treg depletion has proven efficient in impeding tumour growth in preclinical mouse models (Jang et al., 2017; Keenan et al., 2014; Leao et al., 2008; Shevchenko et al., 2013), although a recent report contradicted these findings (Zhang et al., 2020). Here we confirmed that punctual Treg depletion provides immunity against pancreatic tumour growth in orthotopic tumours, and additionally demonstrate the existence of histological alterations to the non-tumoral exocrine parenchyma after tumour regression, which are reminiscent of pancreatitis-induced ADM. These results suggest that the protective effect of pancreatic Tregs on the exocrine parenchyma may be compromised in the context of Treg-targeted therapy of pancreatic cancer, despite normal or elevated pancreatic Treg numbers. Interestingly, recent data suggest that checkpoint inhibitors-induced pancreatitis incidence is increased in patients treated with anti-CTLA-4 antibodies compared to those receiving anti-PD-1 inhibitors, although this meta-analysis did not include any pancreatic tumours (George et al., 2019). Thus, our data argue for the enhanced monitoring of pancreatic adverse events in the context of therapies targeting Treg numbers or function.

Our work illustrates the profound impact stemming from the dualistic nature of trTregs. Hence, understanding the molecular mechanisms underpinning their roles in tissue homeostasis restoration versus immune regulation remains a major challenge. In the context of cancer, selectively targeting the Treg harmful immune-suppressive effects whilst preserving tissue repair capabilities will be of significant importance for future research.

## Material and methods

### Mice

*Foxp3^DTR^* (Kim et al., 2007)(JAX #016958) and *Foxp3^YFP-Cre^*(Rubtsov et al., 2008)(JAX #016959) mice were bought from JAX, and were maintained in the Cancer Research UK – Cambridge Institute (CRUK-CI) animal facility, under specific-pathogen-free conditions along with *Metrnl^−/−^*, *Il33^Cit/Cit^*(Hardman et al., 2013)(provided by Prof. McKenzie), *Nr4a1^Kathushka-CreERT2^ Rosa26^LSL-YFP^* (Takahashi et al., 2023)(provided by Dr. Thaventhiran), *Areg^−/−^* (Zaiss et al., 2013)(provided by Prof. Clatworthy) and *Ptf1a^Cre/+^ Kras^LSL-G12D/+^* (provided by Prof. Brindle) mice, all in the C57Bl/6 background. Wild-type C57Bl/6J mice were purchased from Charles River. Animal work was conducted under project licences PD7484FB9 or PF993249 at CRUK-CI (with approval from CRUK-CI Animal Welfare Ethical Review Body) all in accordance with Home Office regulation. Mice were sex and age matched whenever possible, and most mice were used at 10-16 weeks of age.

### *Metrnl^−/−^* mouse generation

A phosphorothioate-modified sgRNA (5’ CAGAGCCTGCGGAGCAGCGC 3’) was designed against exon 2 of *Metrnl* (ENSMUST00000036742.13). C57Bl/6J one-cell zygotes were electroporated (Nepa21, Sonidel) using the 5 mm electrode (CUY505P5) with 333 ng/µl Cas9 (TrueCut spCas9 protein V2, Invitrogen) and 75 ng/µl guide RNA (Synthego). Two-cell stage embryos were surgically transferred into the oviducts of pseudo-pregnant F1 CBAxC57BL/6J surrogate females (Charles River), using 15 embryos per oviduct. Editing of the *Metrnl* allele was confirmed by Sanger sequencing, and one founder male was used for breeding to a C57Bl/6J female (Charles River) for 2 generations before using animals for experimental purposes.

### *In vivo* procedures

rmIL-33 (0.2 µg, Biolegend) and diphtheria toxin (10 ng/g in AP or 25 ng/g in orthotopic PDAC, Sigma) were administered by i.p. injection in 200 µl of PBS. Labelling of circulating leukocytes to evaluate tissue-residency was achieved by i.v. injection of 3 µg anti-CD45.2-PE (clone 104, Biolegend) antibody 3 min prior to sacrifice. For TCR fate-mapping analyses, AgRSR mice were injected i.p. with 2mg Tamoxifen (Sigma) dissolved in sunflower oil, followed by sacrifice one week later. For AP experiments, mice were given s.c. analgesia before 6 hourly injections of 75 µg/kg Caerulein (Sigma) or endotoxin-free PBS. Anti-MHCII mAb (500 µg, clone M5/114, BioXCell) and rat IgG2b isotype control (500 µg, clone LTF-2, BioXCell) were administered by i.v. injection in 100 µl of PBS one day prior to AP induction. For orthotopic PDAC allografts, mice were anaesthetized with isoflurane and an incision was made in the left abdominal area to expose the pancreas. 10 μl of 1:1 PBS and matrigel cell suspension containing 25 000 KPCY-derived 2838c3 cells (Li et al., 2018b) (kindly gifted by Ben Stanger, University of Pennsylvania) was injected into the tail of the pancreas. The peritoneal membrane was sutured with absorbable Vicryl Rapide suture (Ethicon, W9913) and the skin was closed with tissue glue (GLUture, Zoetis) and wound clips (FST, 12022-09). Mice were given pre- and post-surgical analgesia and wound clips were removed one week post-surgery. For survival studies, the humane endpoint was determined using a combination of clinical signs (e.g. inactivity, piloerection and a hunched posture). All survival studies were terminated at day 90 post-implantation, at which point all mice were culled.

### Human samples

Primary pancreas tissue was obtained by the Cambridge Biorepository of Translational Medicine (CBTM) from deceased organ donors from whom multiple organs were being retrieved for transplantation. Pancreas samples were taken via two routes: from donors during the organ retrieval operation (in which organs other than the pancreas were taken for transplant) or from pancreata which were initially removed for organ transplantation but were subsequently declined and allocated for research. Tissue samples were placed in cold University of Wisconsin organ preservation solution prior to transportation to the laboratory. Donor tissue was taken after obtaining written informed consent from the donor’s family for studies approved by the NRES Committee East of England, Cambridge South for the Department of Surgery, University of Cambridge, REC reference; 15/EE/0152 and the NRES Committee East of England - Cambridgeshire and Hertfordshire Research Ethics Committee for the Department of Surgery, University of Cambridge, REC reference; 16/EE/0227.

### Histology

Tissues were fixed in 10% neutral buffered formalin and embedded in paraffin before sectioning into 4-μm slices. Sample embedding, sectioning and staining were conducted by the CRUK-CI Histology Core. Sections were stained on a Leica’s automated Bond-III platform for mouse CK19 (clone TROMA-III, DSHB), mouse Foxp3 (clone FJK-16s, eBiosciences), or human FOXP3 (clone 236A/E7, abcam), with antigen retrieval using Proteinase K, Tris-EDTA and Sodium Citrate, respectively. ADM area was quantified on H&E sections using Halo Software (Indica Labs).

### Tissue digestion and single cell suspensions

Pancreata were weighed, then mechanically dissociated, followed by digest in 5 ml of HBSS containing collagenase I (375 U/ml), DNase I (0.15 mg/ml) and Soybean Trypsin inhibitor (Sigma, 0.05 mg/ml) for 30 minutes at 37°C on a shaker (220 rpm), followed by dissociation on a syringe and needle, filtration through a 70 μm strainer to exclude Langerhans islets, and red blood cell lysis. pLN were digested in 0.5 ml of HBSS containing collagenase I (375 U/ml) and DNase I (0.15 mg/ml) for 45 min at 37°C on a shaker (1100 rpm).

### Flow cytometry

For intracellular Areg detection, single cells were stimulated with a cytokine stimulation cocktail containing protein transport inhibitors (Thermo Fisher Scientific) for 2 hours at 37C in RPMI supplemented with 10%FCS, 10 µg/ml Soybean Trypsin inhibitor (Sigma) and 10 µM marimastat (Sigma). For Foxp3 and YFP co-staining we followed the protocol described by (Heinen et al., 2014) with intracellular staining performed overnight. Single cells were incubated with anti-mouse CD16/32 (Thermo Fisher) to block Fc receptors. For intracellular staining we used the Foxp3/Transcription Factor Kit (Thermo Fisher). Data was acquired on a BD Fortessa or Symphony instrument. Cells were quantified using CountBright beads. Data was analysed using FlowJo X (Tree Star). The following antibodies were used in this study, with clones, venders, and fluorochrome as indicated: CD45 (30-F11, Biolegend, BV510), CD4 (RM4-5, eBioscience, AF700), CD8 (53-6.7, eBioscience, PerCP-eFluor710), CD3 (145-2C11, eBioscience, PE-Cy7), CD127 (SB/199, BD, PE-CF594), NK1.1 (PK136, BD, BUV395), KLRG1 (2F1, Biolegend, BV605 or eBioscience, PerCP-eFluor710), ST2 (RMST2-2, eBioscience, PE or SB436), TCRβ (H57-597, Biolegend, PE), Gata3 (TWAJ, eBioscience, eFluor660), Foxp3 (FJK-16s, eBioscience, AF488), Areg (polyclonal, R&D Systems, biotin), CD140a (APA5, Biolegend, BV421), CD31 (390, Biolegend, BV605), EpCAM (G8.8, Biolegend, BV711), CD90.2 (30-H12, Biolegend, AF700), CD133 (315-2C11, Biolegend, PE-Dazzle594), Podoplanin (8.1.1, Biolegend, PE-Cy7), Ki67 (B56, BD Biosciences, BUV395). Lineage cocktail contained CD5 (53-7.3), CD19 (1D3), CD11b (M1/70), CD11c (N418), FcεR1α (MAR-1), F4/80 (BM8), Ly-6C/G (Rb6-8C5), and Ter119 (TER-119), all on eFluor450 (eBioscience). Dead cells were excluded with the fixable viability dye UV455 or eFluor780 (eBioscience).

### qPCR

Single cell suspensions from pancreata were used to isolate acinar cells (Live CD45^−^ EpCAM^+^CD133^lo^SSC^hi^), ductal cells (Live CD45^−^EpCAM^+^CD133^+^SSC^lo^) and fibroblasts (Live CD45^−^EpCAM^−^CD90^+^Podoplanin^+^) by flow cytometry using a FACS Aria instrument. RNA was extracted using Tri-Reagent (Sigma) followed by 1-bromo-3-chloropropane extraction and clean-up on RNeasy columns. RNA from total pancreas lysates was prepared as followed. After sacrifice, pancreata were perfused with 500 µl RNA Later (Ambion), and snap frozen in liquid nitrogen. Upon thawing, tissue was directly transferred into CK28 tubes (Precellys) filled up with 5 ml Tri-Reagent (Sigma), and homogenized for 1*30sec at 7200 rpm using a Precellys homogeniser. RNA extraction from homogenates was performed following the Tri-Reagent protocol, with the addition of an isopropanol washing step before ethanol precipitation. RNA quality was assessed using the Agilent Tapestation. Treg cells (Live CD45^+^CD3^+^CD8^−^ CD4^+^TCRβ^+^Foxp3^YFP+^), Tconv cells (Live CD45^+^CD3^+^CD8^−^CD4^+^TCRβ^+^Foxp3^YFP-^) and ILC2 cells (Live CD45^+^CD3^−^CD8^−^Lin^−^CD127^+^) were isolated by flow cytometry from single cell suspensions prepared from pancreata of IL-33-treated *Foxp3^YFP^* mice using a FACS Aria instrument. RNA was converted into cDNA using the High-Capacity RNA-to-cDNA Kit (Thermo Scientific), followed by qPCR using the Takyon Low Rox Probe Master mix dTTP Blue (Eurogentec) and primers/probe pre-designed assays (Integrated DNA Technologies): *Hprt* (Mm.PT.39a.22214828), *Gusb* (Mm.PT.39a.22214848), *Cpa1* (Mm.PT.58.15958828), *Krt19* (Mm.PT.58.7322803), *Pdgfra* (Mm.PT.56a.5639577), *Il33* (Mm.PT.58.12022572), *Areg* (Mm.PT.58.31037760), *Metrnl* (Mm.PT.58.11823014), *Calca* (Mm.PT.58.23694914).

### Acinar cell line culture

The 266-6 pancreatic acinar cell line (CRL-2151) was bought at the ATCC, and cultured in DMEM supplemented with 10% FCS, 100 U/ml penicillin (Gibco), and 100 μg/ml streptomycin (Gibco) at 37°C in a humidified, 5% CO_2_ incubator. Cells were transduced with the Incucyte Nuclight Red (mKate2) Lentivirus reagent (Sartorius) according to the manufacturer’s instructions, and stably selected under 1 µg/ml puromycin (Sigma). Proliferation assay in the presence of rmAreg (Biolegend) was carried out in the Incucyte live imaging system.

### Acinar cell and T cell co-culture

On day -1, 1250 266-6-mKate cells were seeded in 96F plate in DMEM media supplemented with 10% FCS, 100 U/ml penicillin (Gibco), and 100 μg/ml streptomycin (Gibco) (Supplementary Figure 5G). On day 0, Treg cells (Live CD45^+^CD3^+^CD8^−^ CD4^+^TCRβ^+^Foxp3^YFP+^) and Tconv cells (Live CD45^+^CD3^+^CD8^−^CD4^+^TCRβ^+^Foxp3^YFP-^) were isolated by flow cytometry from single cell suspensions prepared from pancreata of IL-33-treated *Foxp3^YFP^* mice using a FACS Aria instrument. 3000 purified T cells in RPMI supplemented with 2.5%FCS were added to pre-washed 266-6 mKate cells, along with rmIL-33 (20 ng/ml, Biolegend), rhIL-2 (50 ng/ml, Peprotech), and Dynabeads Mouse T-Activator CD3/CD28 (2 beads:1 cell ratio, Themo Scientific), followed by incubation in the Incucyte live imaging system for measurement of 266.6-mKate proliferation.

### Microscopy

Whole pancreas explants from *Il33^Cit/+^* mice were imaged on an inverted laser-scanning confocal microscope (Stellaris 8, Leica Microsystems) with an attached pulsed Coherent Chameleon Ultra II laser, using a 25X 0.95NA water immersion objective lens. Citrine was excited at a single-photon excitation wavelength of 900nm, and collagen SHG at a 2-photon excitation wavelength of 800nm. Immunofluorescence imaging was performed on FFPE sections by staining with primary antibodies against podoplanin (8.1.1, BioLegend) and IL-33 (EPR17831, abcam) overnight at 4°C. Secondary antibodies used were an anti-Syrian hamster coupled to AF647 (Stratech) and an anti-rabbit coupled to AF555 (Abcam). Multiplex immunofluorescence imaging was performed on FFPE sections using an Opal multiplex IHC kit (Akoya Biosciences), following the manufacturer’s instructions. Briefly, tissue sections were subjected to antigen retrieval using 10 mM sodium citrate (pH 6) buffer, followed by blocking with the blocking/antibody diluent for 20 min at RT. Subsequently, sections were incubated with primary antibodies against Foxp3 (FJK-16s, Thermo Scientific), IL-33 EPR17831, Abcam), and podoplanin (8.1.1, Biolegend) during respective rounds of iterative staining. After washing, HRP-conjugated secondary antibody was then incubated for 15 minutes (anti-rat and anti-rabbit polymer-HRP) or 30 minutes (anti-Syrian hamster-HRP). Sections were washed prior to addition of Opal reagents (Opal 570, Opal 520, Opal 690) in Opal amplification buffer.

### Bulk RNA-seq

Treg (CD45^+^Lin^−^CD3^+^CD4^+^CD8^−^YFP^+^KLRG1^+^ or KLRG1^−^) cells were sorted from pancreata or pLN of naïve or IL-33-treated *Foxp3^YFP^* reporter mice using a BD Aria II cell sorter. For the naïve mice dataset, cells were sorted directly in a guanidine 6M buffer according to the protocol described by (Wollny et al., 2016). Briefly, cells were sorted in 0.2 ml tubes containing 4 μl per well of 0.2% Triton-X100, 1% 2-mercaptoethanol, 6M guanidine HCl in dH_2_O. Cellular RNA was cleaned with 2.2x volume of RNAclean XP SPRI beads per sample (Beckman Coulter). Dried beads were resuspended with 4 μl of MM1 solution (10 nM dNTP, 10 μM oligo-dT30VN, 2.5% v/v SuperRNAsin) and incubated for 5 minutes, followed by reverse transcription by addition of 5.5 μL of MM2 reaction mix, and subsequent PCR with the KAPA HiFi HotStart Ready Mix (Roche) and ISO SMART primer as per the standard protocol. Following PCR, products were cleaned-up with AMPure XP beads (Beckman Coulter). For the IL-33-treated mice dataset, cells were sorted into PBS-2% FCS and RNA was extracted according to (Halim et al., 2018), except for 1-bromo-3-chloropropane used instead of chloroform. 1 ng RNA was used as input to generate cDNA with the SMART-Seq v4 Ultra Low Input RNA Kit (Takara), according to manufacturer’s instructions. For both datasets, libraries were generated with the Nextera XT kit (Illumina) using 500 pg of cDNA as per the manufacturer’s instructions, and sequenced (paired end 150nt) on an Illumina NovaSeq 6000 system (Illumina).

### Bulk RNA-seq analysis

Raw sequence data quality was assessed using FastQC. Raw reads were trimmed for adapter content using Trimmomatic and aligned to the GRCm38.p6 genome using STAR. Alignment quality was assessed using Picard’s tools. A matrix of counts of reads aligned against genes was generated using the featureCounts programme from the Subread package and the Ensembl release 99 gene annotation GTF file. In the naïve dataset, 2 samples were excluded based on hierarchical clustering analyses. Furthermore, we removed 33 acinar cell-specific genes (based on our pancreas scRNA-seq data, Yip *et al.,* submitted) from both datasets as they were found to be driving variation in the first principal component of the pancreas-derived samples. The list of these genes can be found in Supplementary Table 3. Analysis of differential gene expression was carried out using the DESeq2 package. T-cell receptor clonotypes were extracted from the raw fastq files using MiXCR. Diversity metrics along with abundance analyses were carried out with the Immunarch package in R. Data have been deposited to NCBI GEO under GEO accession number GSE240050.

### Cell Chat analysis

Pancreatic Tregs scRNA-seq data were obtained from (Burton et al., 2023). Pancreatic Tregs with fewer than 5% mitochondrial transcripts were used for downstream analysis. Cells were clustered using standard graph-based clustering methods in Seurat performed on the first 30 principal components, resulting in two clusters (FindClusters resolution of 0.2) that demonstrated differential expression of tissue-residency markers including *Gata3*, *Klrg1*, and *Il1rl1*. Cells belonging to this trTreg cluster were then merged with scRNA-seq transcriptomic data of acinar cells from a time course of pancreatic injury and repair (Yip *et al.,* submitted). Potential receptor-ligand interactions from Treg acting on acinar cells were inferred using the CellChat package. Computation of communication probability was performed using the ‘truncatedMean’ method with trim=0.1.

### Statistics

Analysis for two groups were calculated using an unpaired two-tailed Student’s *t*-test; comparisons of more than two groups were calculated using a one-way analysis of variance (ANOVA) with Tukey post-analysis, except for matched samples where the Holm-Šídák post-analysis was used. When one or more matched values were missing, comparisons were calculated using a mixed-effects analysis with Holm-Šídák post-analysis. All data were analysed using GraphPad Prism 10 (GraphPad Software). Results with p ≤ 0.05 being considered significant (*), p ≤ 0.01 = **, p ≤ 0.001 = ***, p ≤ 0.0001 = ****.

## Acknowledgements

We thank the CRUK-CI research instrumentation, flow cytometry, genomics, bioinformatics, histopathology, microscopy, transgenics and BRU cores for their expertise and help. We are grateful to the donors and their families for the precious gift of tissues, provided through the Cambridge Biorepository for Translational Medicine.

