## Supplemental Figures 1-7 for "Tissue-resident regulatory T cells exert dualistic anti-tumour and pro-repair function in the exocrine pancreas"

### Supplementary material

### Supplementary Figures and Supplementary Figure Legends

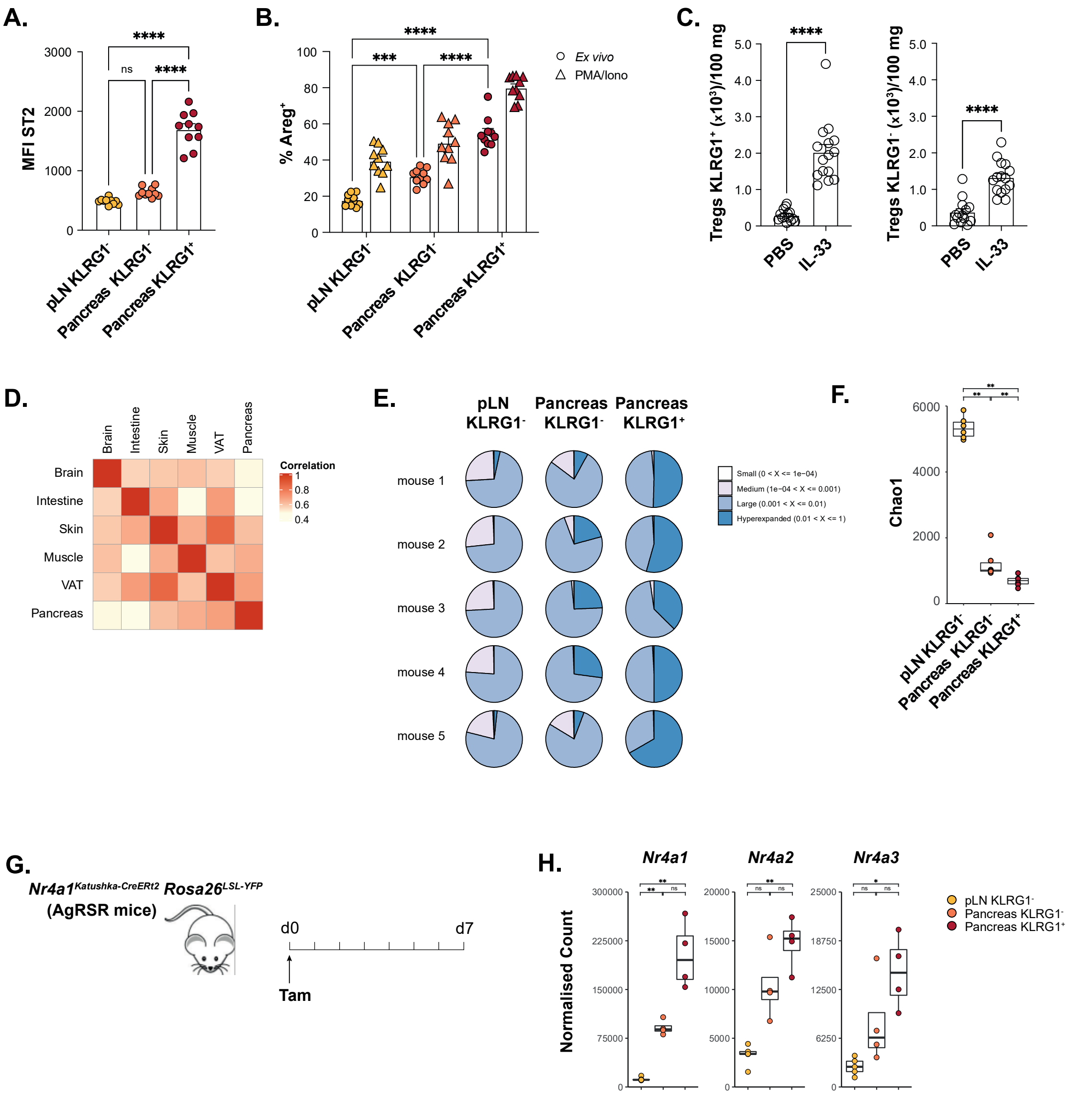


**Supplementary Figure 1.**

Pancreatic KLRG1$^{+/-}$ and pLN KLRG1$^{-}$ Tregs were analysed for ST2 expression by flow cytometry (**A**). Single cell suspensions from pancreas or pLN were re-stimulated *in vitro* with PMA and ionomycin followed by quantification of Areg production by flow cytometry in the indicated populations (**B**). Wild-type mice were dosed i.p. with IL-33 on days 0 and 1, and sacrificed on day 5, followed by quantification of KLRG1$^{+}$ or KLRG1$^{-}$ pancreatic Treg numbers by flow cytometry (**C**). Correlation matrix comparing Treg bulk RNAseq datasets from different tissue origin (**D**). TCR-β diversity analysis on the indicated populations from the bulk RNA-seq dataset obtained from naïve (**E**) or IL-33-treated mice (**F**). Mice of the indicated genotype were treated with tamoxifen and sacrificed 7 days later (**G**). Expression of indicated genes from the pancreatic Treg bulk RNA-seq samples obtained from naïve mice (**H**).

Bar graphs indicate mean (±SEM) and show representative data of >3 independent experiments (n=10 mice, A, B), or pooled data from 3 independent experiments (n=4 to 5 mice per group, C).
* = p ≤ 0.05, ** = p ≤ 0.01, *** = p ≤ 0.001, **** = p ≤ 0.0001, ns = not significant.

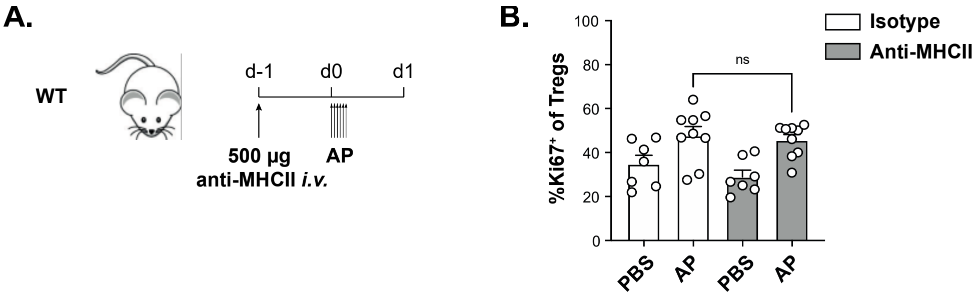


**Supplementary Figure 2.**

Wild-type mice were dosed with anti-MHC-II antibody before induction of AP as indicated, and sacrificed one day later (**A**), followed by quantification of cycling Tregs by flow cytometry (**B**).

Bar graphs indicate mean (±SEM) and show pooled data from 2 independent experiments (n=3 to 5 mice per group, B).

* = p ≤ 0.05, ** = p ≤ 0.01, *** = p ≤ 0.001, **** = p ≤ 0.0001, ns = not significant.

A.

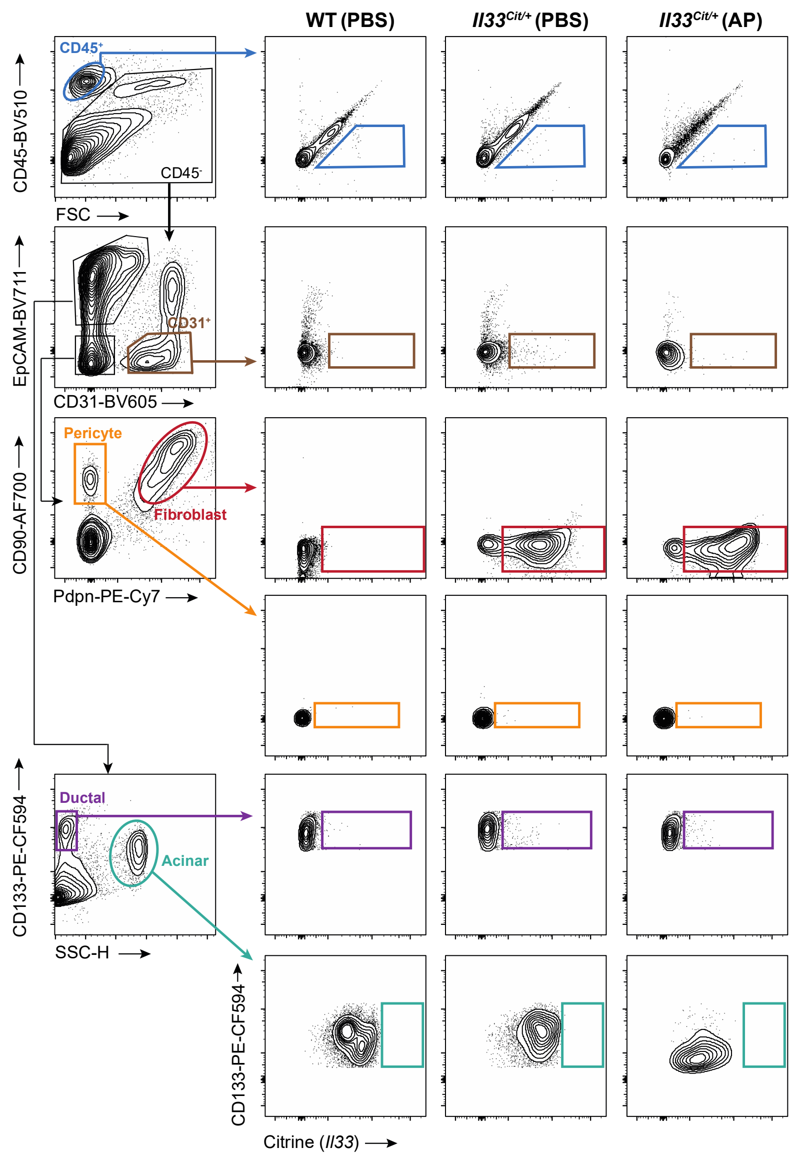
WT (PBS)
Il33Cit/+ (PBS)
Il33Cit/+ (AP)
CD45+
CD45-
CD45-BV510
FSC
EpCAM-BV711
CD31+
CD31-BV605
Pericyte
Fibroblast
CD90-AF700
Pdpn-PE-Cy7
Ductal
Acinar
CD133-PE-CF594
SSC-H
CD133-PE-CF594
Citrine (Il33)

B.

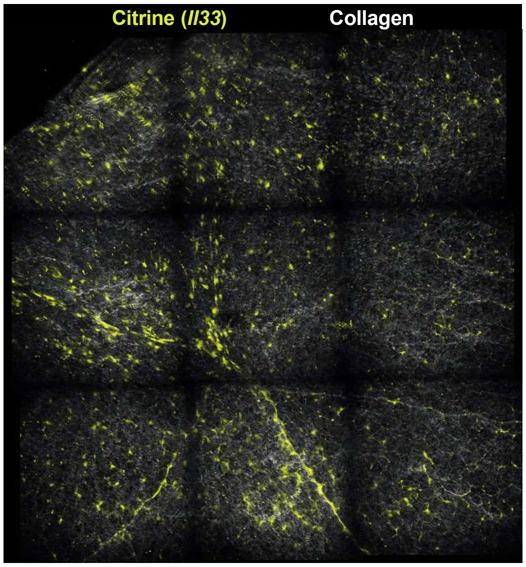
Citrine (Il33)
Collagen

**Supplementary Figure 3.**

Representative gating strategy for the major non-immune cell subsets of the pancreas. Endothelial cells are gated as $CD45^-EpCAM^-CD31^+$, pericytes as $CD45^-EpCAM^-CD31^-CD90^+Pdpn^-$, fibroblasts as $CD45^-EpCAM^-CD31^-CD90^+Pdpn^+$, acinar cells as $CD45^-CD31^-EpCAM^+SSC^{hi}$, and ductal cells as $CD45^-CD31^-EpCAM^+SSC^{lo}CD133^{hi}$ (**A**). Representative regions of 2-photon imaging of pancreas explants from naive *Il33*$^{Cit/+}$ reporter mice (**B**).

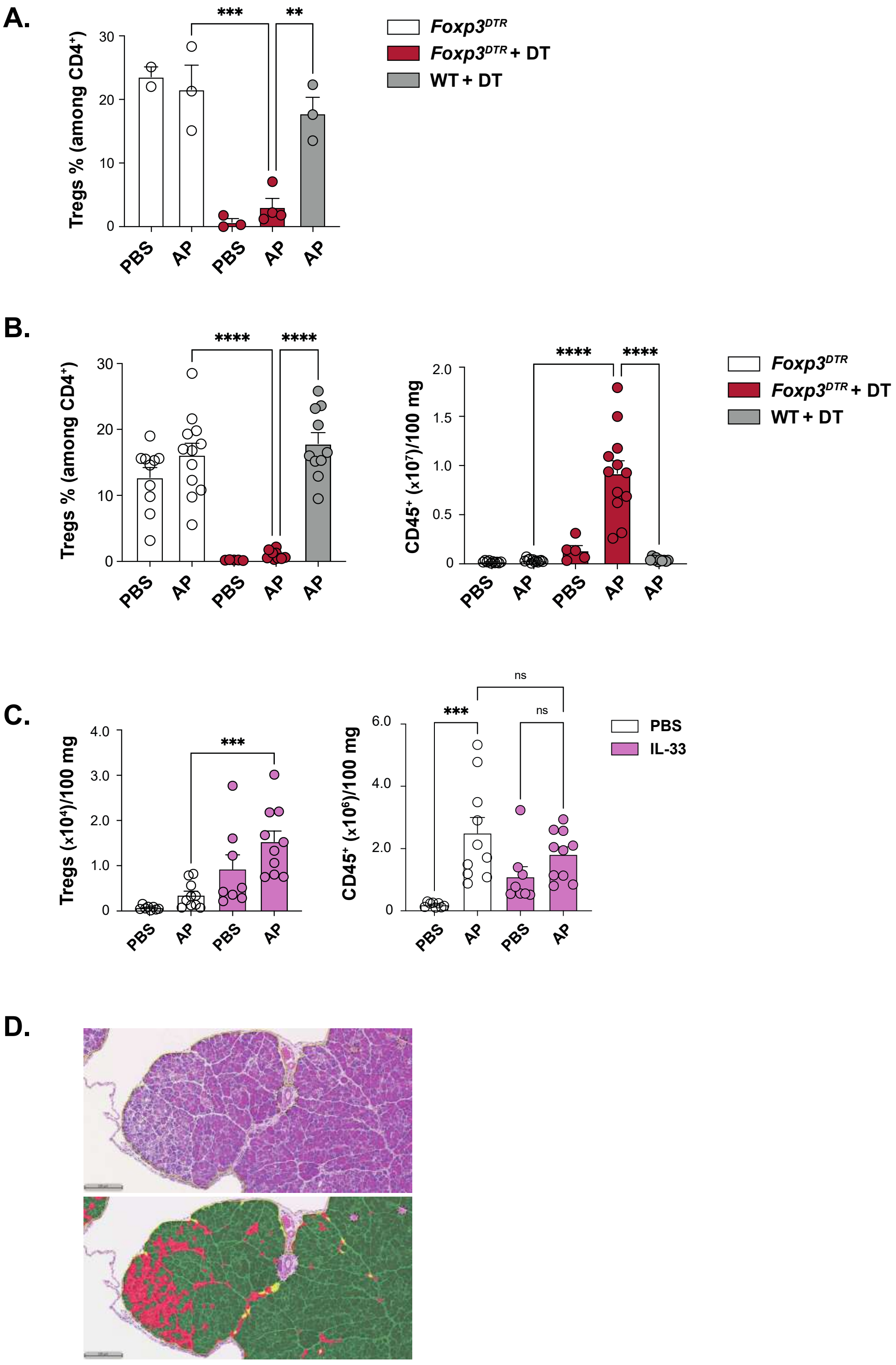


**Supplementary Figure 4.**

Mice of the indicated genotype were treated with diphtheria toxin (DT) before induction of AP and sacrificed one day later, followed by quantification of Treg as a proportion of $CD4^+$ T cells by flow cytometry (**A**). Mice of the indicated genotype were treated with DT after the induction of AP and sacrificed 7 days later, followed by quantification of Treg as a proportion of $CD4^+$ T cells and total $CD45^+$ numbers by flow cytometry (**B**). Wild-type mice were dosed with IL-33 before induction of AP and sacrificed 2 days later, followed by quantification of total Treg (left) and $CD45^+$ (right) numbers by flow cytometry (**C**). Representative classifier mask for ADM scoring on H&E stained FFPE sections with the HALO image analysis software (**D**).

Bar graphs indicate mean (±SEM) and show representative data of 2 independent experiments (n=3 to 4 mice per group, A), or pooled data from 2 (n=4 to 5 mice per group, C) to 3 independent experiments (n=3 to 5 mice per group, B).
* = p ≤ 0.05, ** = p ≤ 0.01, *** = p ≤ 0.001, **** = p ≤ 0.0001, ns = not significant.

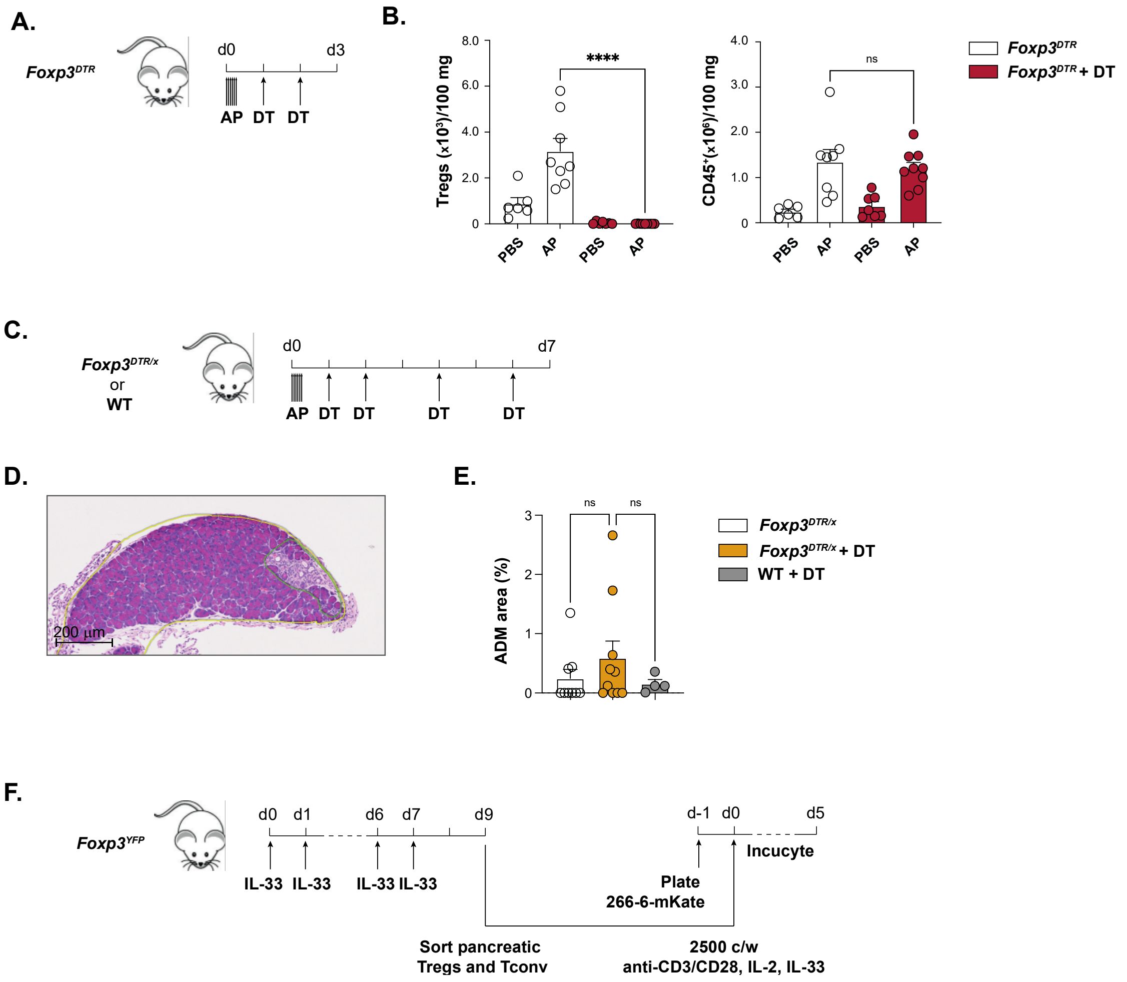


**Supplementary Figure 5.**

Mice of the indicated genotype were treated with DT after the induction of AP and sacrificed 3 days later (**A**), followed by quantification of total Treg (left) and CD45$^+$ (right) numbers by flow cytometry (**B**). Mice of the indicated genotype were treated with DT after the induction of AP and sacrificed 7 days later (**C**), followed by histological examination of residual ADM patches on H&E stained FFPE sections with the HALO image analysis software (**D-E**). Sorted pancreatic Tregs or Tconv cells were co-cultured with 266-6 acinar cells expressing the red fluorescent protein m-Kate in the presence of anti-CD3/anti-CD28 coated beads, IL-2 and IL-33. Acinar cell proliferation was measured in the Incucyte live imaging system (**F**).

Bar graphs indicate mean (±SEM) and show pooled data from 2 independent experiments (n=3 to 5 mice per group, B, E).

* = p ≤ 0.05, ** = p ≤ 0.01, *** = p ≤ 0.001, **** = p ≤ 0.0001, ns = not significant.

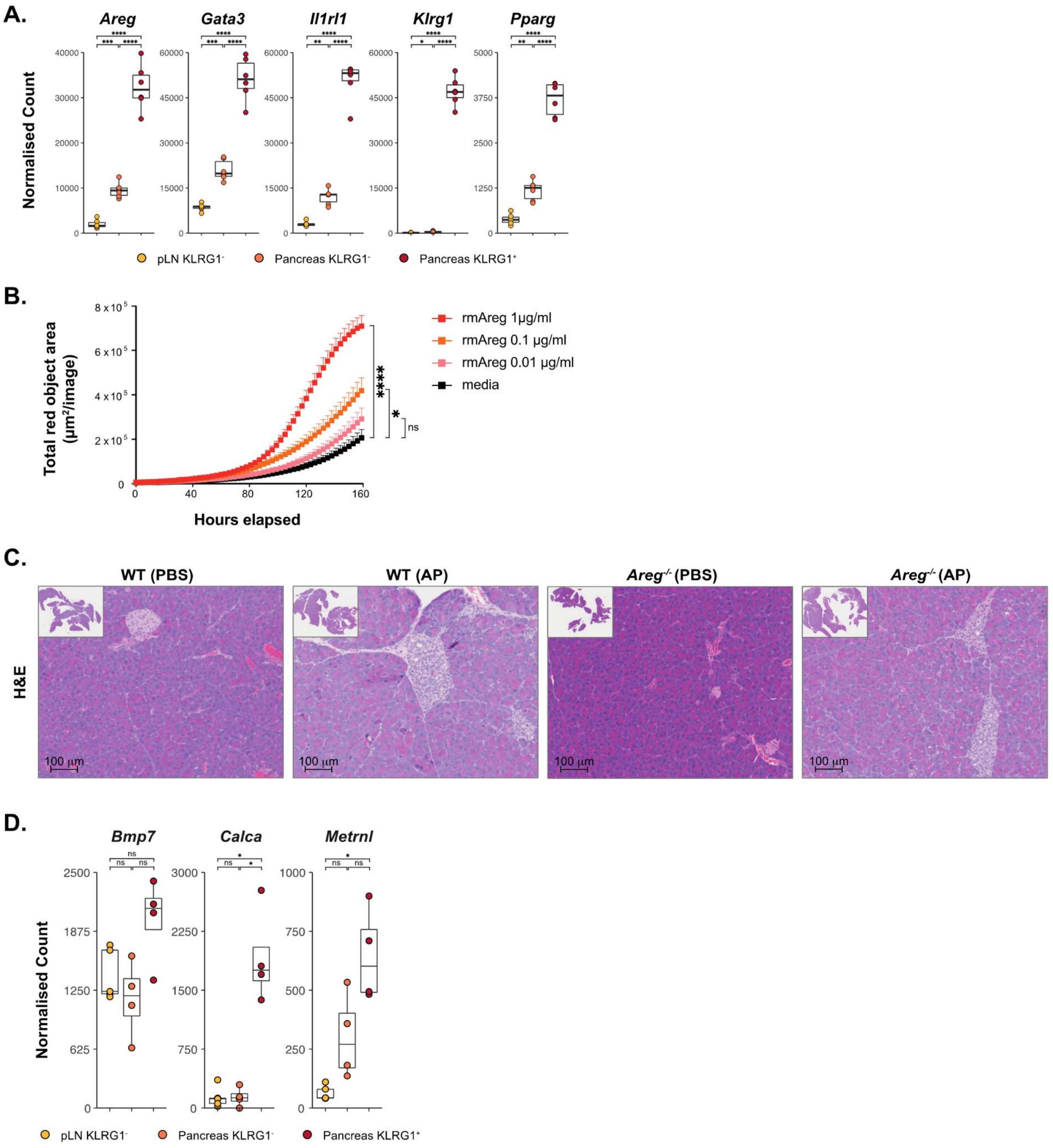


**Supplementary Figure 6.**

Expression of indicated genes from the pancreatic Treg RNA-seq samples obtained from mice treated with IL-33 (**A**). Proliferation of 266-6 acinar cells expressing the red fluorescent protein m-Kate were in the presence of indicated concentrations of rmAreg was measured in the Incucyte live imaging system (**B**). Mice of the indicated genotype were sacrificed 7 days after induction of AP, followed by histological examination of H&E sections (**C**). Expression of indicated genes from the pancreatic Treg RNA-seq samples obtained from naïve mice (**D**).

Bar graphs indicate mean (±SEM) and show representative data of 1 independent experiments (n=4 to 6 mice per group, A, D). Growth curve shows data representative of 3 independent experiments.

* = $p \le 0.05$, ** = $p \le 0.01$, *** = $p \le 0.001$, **** = $p \le 0.0001$, ns = not significant.

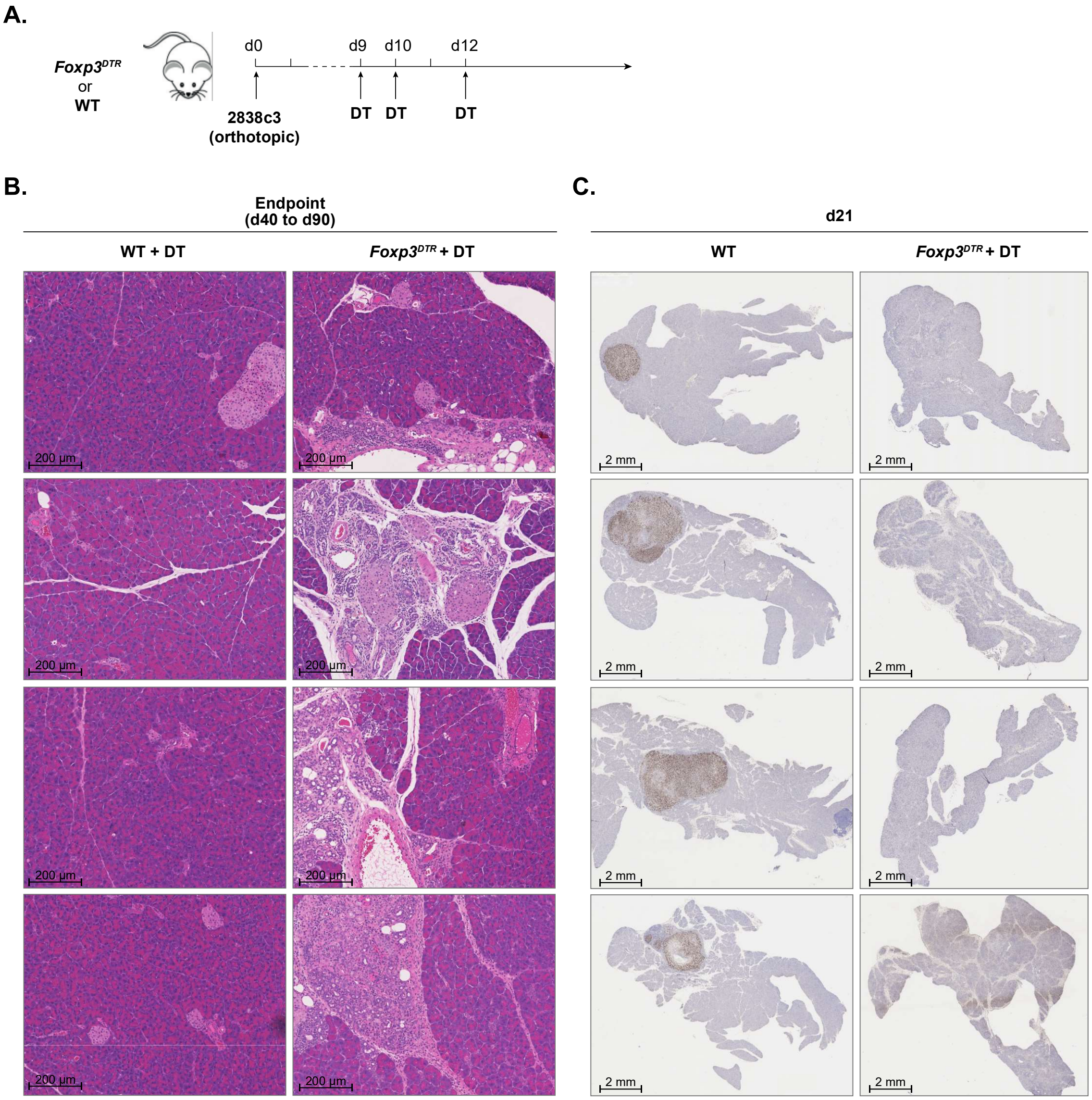


**Supplementary Figure 7. Anti-tumour efficacy of Treg depletion in an orthotopic PDAC model is associated with alterations to the exocrine parenchyma.**

2838c3 cells were implanted orthotopically in the pancreas of mice of the indicated genotype, followed by treatment with DT as indicated (**A**). Representative H&E sections of mice treated as in (A) and sacrificed at the study humane endpoints (**B**). CK19 staining of FFPE sections from mice treated as in (A) and sacrificed at 21 days post-implantation (**C**).
